# The niche is not the range: Dispersal and persistence shape mismatches between ecological niches and geographic distributions of plants

**DOI:** 10.1101/526251

**Authors:** Jörn Pagel, Martina Treurnicht, William J. Bond, Tineke Kraaij, Henning Nottebrock, AnneLise Schutte-Vlok, Jeanne Tonnabel, Karen J. Esler, Frank M. Schurr

**Affiliations:** Institute of Landscape and Plant Ecology, University of Hohenheim, August-von-Hartmann-Str. 3, 70599 Stuttgart, Germany; Department of Conservation Ecology and Entomology, Stellenbosch University, Private Bag X1, Matieland 7602, South Africa; South African Environmental Observation Network (SAEON), Private Bag X7, Claremont 7735, South Africa; Department of Biological Sciences, University of Cape Town, Rondebosch 7701, South Africa; School of Natural Resource Management, Nelson Mandela University, George 6529, South Africa; Department of Biology and Microbiology, SD AES, South Dakota State University, Brookings, South Dakota 57005, USA; Plant Ecology, University of Bayreuth, Universitätsstrasse 30, 95447 Bayreuth, Germany; Scientific Services, CapeNature, Private Bag X658, Oudtshoorn 6620, South Africa; Department of Ecology and Evolution, Le Biophore, Quartier UNIL-Sorge, University of Lausanne, 1015 Lausanne, Switzerland

## Abstract

The ecological niche of a species describes the variation in population growth rates along environmental gradients that drives geographic range dynamics. Niches are thus central for understanding and forecasting species’ geographic distributions. However, theory predicts that migration limitation, source-sink dynamics and time-lagged local extinction can cause mismatches between niches and geographic distributions. It is still unclear how relevant these niche-distribution mismatches are for biodiversity dynamics and how they depend on species life history traits. This is mainly due to a lack of the comprehensive, range-wide demographic data needed to directly infer ecological niches for multiple species. Here we quantify niches from extensive demographic measurements along environmental gradients across the geographic ranges of 26 plant species (Proteaceae; South Africa). We then test whether life history explains variation in species’ niches and niche-distribution mismatches. Niches are generally wider for species with high seed dispersal or persistence abilities. Life history traits also explain the considerable interspecific variation in niche-distribution mismatches: poorer dispersers are absent from larger parts of their potential geographic ranges, whereas species with higher persistence ability more frequently occupy environments outside their ecological niche. Our study thus identifies major demographic and functional determinants of species’ niches and geographic distributions. It highlights that the inference of ecological niches from geographical distributions is most problematic for poorly dispersed and highly persistent species. We conclude that the direct quantification of ecological niches from demographic responses to environmental variation is a crucial step towards a better predictive understanding of biodiversity dynamics under environmental change.

---

In 1957, George Evelyn Hutchinson introduced his seminal concept of a species’ ecological niche (1). The Hutchinsonian niche is defined as the set of environmental conditions for which demographic rates result in a positive intrinsic population growth rate and thus permit a species to form self-perpetuating populations (2, 3). This niche concept has received much attention as a theoretical foundation for explaining the geographic distributions of species and forecasting range shifts under environmental change. However, this use of the Hutchinsonian niche concept has been critically revisited in recent years (4, 5). Theoretical models of range dynamics predict that the geographic distribution of a species does not necessarily match the geographic projection of the species’ niche. This is because geographic distributions are structured by extinction and colonization events that arise from a dynamic interplay of spatial variation in demographic rates, local population persistence and dispersal (6). Specifically, migration limitation can prevent a species from colonizing parts of its potentially suitable range, source-sink dynamics can sustain populations by immigration even if local population growth rates are negative, and time-lagged local extinction can cause species to occur in locations that became unsuitable due to environmental change (4-6). Strong mismatches between niches and geographic distributions can severely bias niche estimates and forecasts of species distribution models (7, 8), which are widely used in global change-biodiversity assessments (9). However, the extent of these niche-distribution mismatches is poorly known, mainly because of a lack of the comprehensive, range-wide demographic data needed to directly infer ecological niches (5, 10-12). Demography-based niches were so far only quantified for a few single species (13-15), which precluded comparative analyses. More than 60 years after Hutchinson introduced his niche concept, it is thus unclear how relevant niche-distribution mismatches are in the real world and how they depend on the dispersal and persistence ability of species.

Here we quantify niches from extensive demographic data for 26 closely related plant species to analyse how niche sizes, geographic range sizes and, finally, niche-distribution mismatches depend on species’ life history traits. Our study species are shrubs of the Proteaceae family endemic to the Cape Floristic Region, a global biodiversity hotspot (16). All study species are serotinous: they store their seeds over multiple years in a canopy seedbank until fire triggers seed release, wind-driven seed dispersal and subsequent establishment of new recruits (17). Fire is also the predominant cause of mortality of established adults (18). This fire-linked life cycle allows the efficient measurement of key demographic rates in single visits to each population (19) and enabled us to collect data that are informative of variation in population growth rates across each species’ geographic range.

## Results and Discussion

We analysed a total of 3,617 population-level records of fecundity, recruitment and adult fire survival (Tab. S1). The analyses used hierarchical demographic response models that describe for each key demographic rate the species-specific response curves to variation in climatic-edaphic conditions (minimum winter temperature, maximum summer temperature, summer aridity, soil fertility), fire return intervals and intraspecific density (Fig. 1a; see Fig. S1 for all study species). The fitted response curves explained much variation in long-term fecundity (Nagelkerke’s R^2^_N_ 0.29–0.91 across species, mean = 0.58; Tab. S2), recruitment (R^2^_N_ = 0.05–0.84, mean = 0.42) and adult fire survival (R^2^_N_ = 0.41–0.83, mean = 0.62). The response curves of all three demographic rates were then integrated in models of density-dependent local population dynamics that predict variation of intrinsic (low-density) population growth rates (*r*_0_) along environmental gradients. This delimits each species’ niche as a hypervolume in environmental space for which *r*_0_ is positive (Fig. 1b). Within species, different demographic rates often respond similarly to the same environmental variable. However, we also found some opposing responses, in particular between fecundity and recruitment rates (for an example see responses to aridity in Fig.1a). Such opposing responses indicate demographic compensation that can broaden species’ environmental niches and buffer effects of environmental change (15, 20-21).

**Fig. 1.**
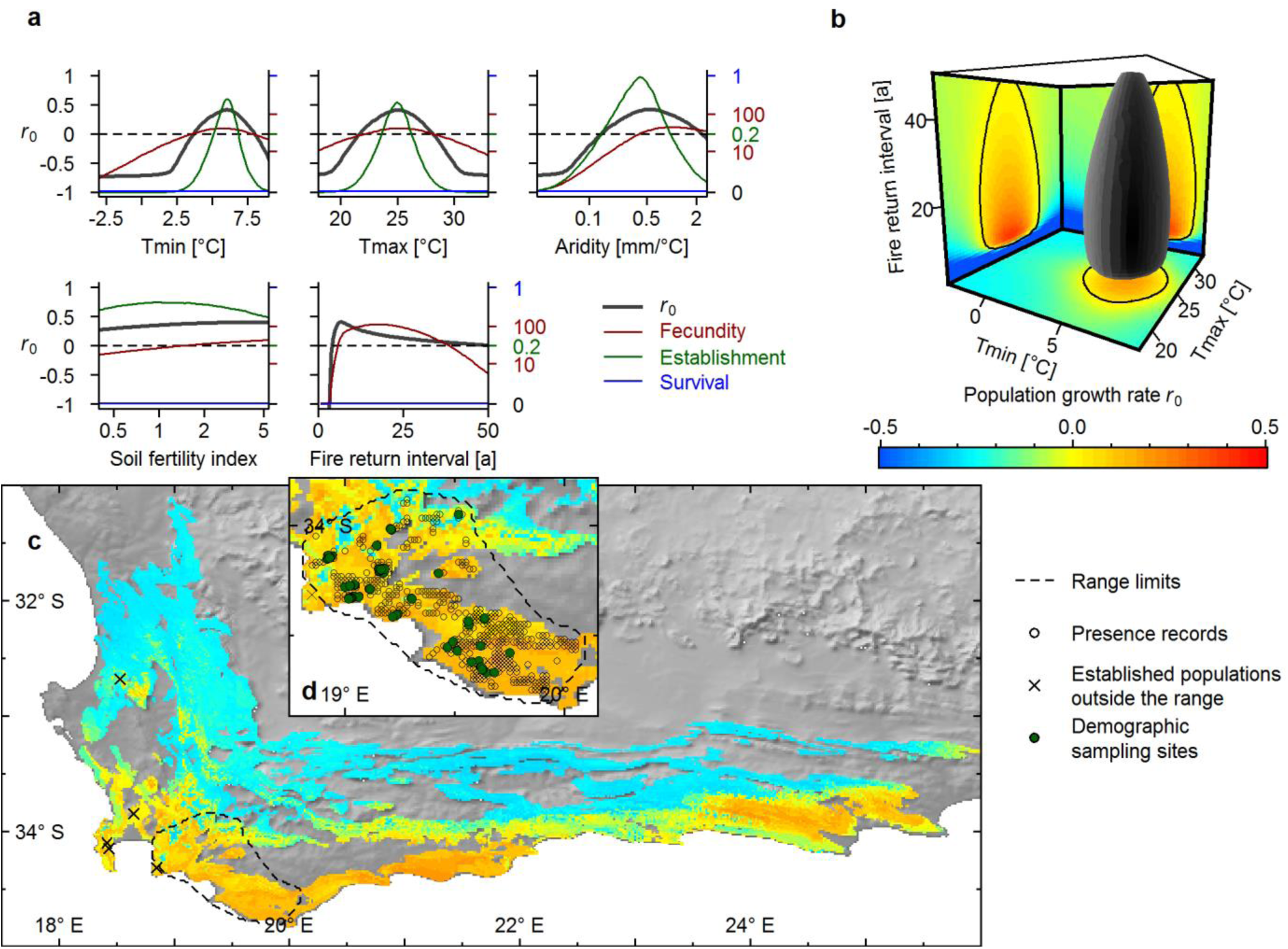
The Hutchinsonian niche and geographic distribution of *Protea longifolia*. (a) Responses of key demographic rates and the resulting annual intrinsic population growth rate (r_0_) to variation in minimum winter temperature (T_min_), maximum summer temperature (T_max_), indices of summer aridity and soil fertility, and fire return interval. (b) The niche hypervolume (grey) in a 3-d environmental sub-space delimits the conditions for which r_0_ > 0. The marginal 2-d heat maps show the predicted r_0_ when all other niche axes are set to their respective optima. (c) Geographic projection of r_0_ across the Fynbos biome (coloured areas) in comparison to the natural geographic range (dashed line) and to populations established outside the natural range (crosses). (d) Enlarged map showing presence records of natural populations (open circles) and demographic sampling sites (green circles). Model predictions in all subplots are the medians of the respective Bayesian posterior distributions.

### Relationship between life history trait effects and niche sizes

We first examined how the size of the estimated niches is related to the dispersal and persistence abilities of species. Niche sizes were quantified separately for the four-dimensional ‘environmental niche’ defined by spatially varying long-term averages of climatic-edaphic conditions and for the ‘disturbance niche’ defined by fire return intervals that also show strong temporal variation in any given location (22). Dispersal ability was quantified from species-specific parameterizations of a trait-based, mechanistic model of wind-driven seed dispersal (23). Persistence ability was characterized by resprouting as a key functional trait: Some of the study species (‘resprouters’, n = 7) possess fire-protected meristems from which individuals can resprout and are thus more likely to survive fire than individuals of species lacking this trait (‘nonsprouters’, n = 19) (18, 19). Since resprouter populations do not exclusively rely on successful reproduction in each fire cycle, they are expected to be less vulnerable to short fire return intervals that prevent the build-up of canopy seed banks (18). We indeed found that resprouting ability had a clear positive effect on disturbance niche size (Fig. 2a), whereas dispersal ability had no effect (Tab. S3). In contrast, environmental niche size showed a strong positive relationship with dispersal ability (Fig. 2b). This finding is consistent with a scenario of correlational selection on niche size and dispersal, where narrower environmental niches select for lower dispersal distances and vice versa (24, 25).

**Fig. 2.**
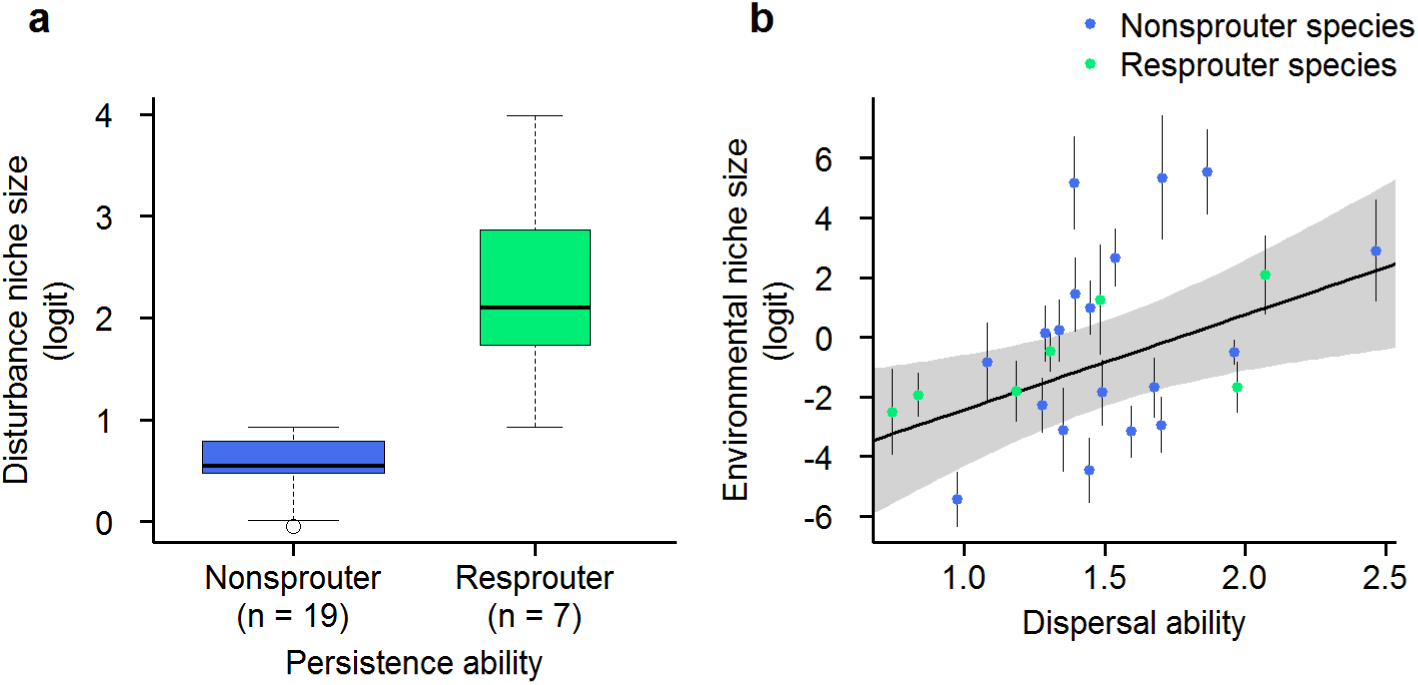
Life history trait effects on niche sizes. (a) Effect of persistence ability on disturbance niche size. (b) Effect of dispersal ability on environmental niche size (points: posterior means, bars: posterior standard deviations). The line shows the estimated linear regression (posterior means, 90% credibility interval as shaded areas, slope = 1.22, *p* = 0.016).

### Mismatches between demographic niches and geographic distributions

For each species, we projected niches from environmental space into geographic space (Fig. 1c; see Fig. S2 for all study species) and then compared this potential geographic range (the region of demographic suitability where predicted *r*_0_ > 0) to independent and extensive distribution records (17). Several species showed a remarkably strong agreement between potential ranges and observed geographic distributions (AUC values >0.8 for ten of the 26 study species). However, there also was a high variation in this agreement across species (AUC 0.55-0.97, mean = 0.77; Tab. S2), indicating interspecific variation in mismatches between demographic suitability and geographic distributions. Since processes that can generate these mismatches are expected to act on different spatial scales (26), we further analysed the relationship between demographic suitability and species occurrence separately at large and small spatial scales.

On large spatial scales, dispersal limitation can cause an incomplete filling of potential ranges, since species are unable to reach suitable areas (4-6). We thus tested for a positive relationship between dispersal ability and range filling. Range filling was measured as the proportion of the potential range that is covered by a species’ geographic range (the alpha-convex hull encompassing all natural occurrences, Fig. 1d). As expected, range filling strongly increased with dispersal ability (Fig. 3a). Hence, good dispersers not only have larger environmental niches (Fig. 2b) and thus tend to have larger potential geographic ranges but they also fill more of their potential ranges, so that both factors add up to explain the larger geographic ranges of good dispersers (Tab. S3). In contrast, persistence ability had no effect on range filling. Absence from suitable areas could of course also indicate that occurrence is limited by environmental factors not considered in our analyses. Ideally, transplant experiments could be used to test model predictions of suitable areas outside the range (27). Such large-scale transplant experiments do, however, pose substantial ethical and logistic problems (28). Instead, we made use of the fact that transplantation by humans (notably flower producers) caused most of our study species to form naturalized populations in natural ecosystems outside their native geographic range (Fig. 1c) (17). When evaluating our model extrapolations for these naturalized populations, we found that the predicted *r*_0_ was positive for an average of 80% populations per species (Table S4). This quasi-experimental evidence suggests that the demographic niche models capture key factors limiting the geographic distributions of our study species.

**Fig. 3.**
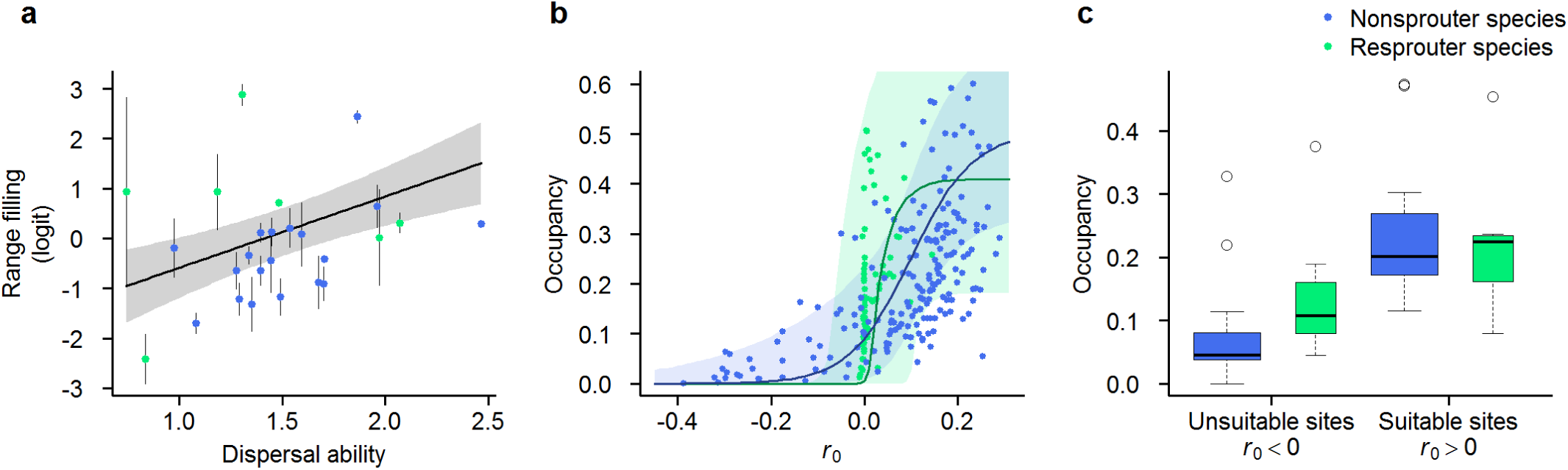
Life history trait effects on the mismatch between niches and geographic distributions. (a) Effect of dispersal ability on range filling (points: posterior means, bars: posterior standard deviations). The line shows the estimated linear regression (posterior means, 90% credibility interval as shaded areas, slope = 0.55, p = 0.022). (b) Relationship between demographic suitability (predicted *r*_0_) and occupancy within the range. Points show the mean occupancy in sites that were binned according to deciles of predicted *r*_0_ (i.e. ten points per species). The lines show average predictions of this relationship for species with different persistence ability (posterior means, 90% credibility interval as shaded areas). (c) Variation in species’ mean occupancy of sites within their ranges that are predicted to be unsuitable (*r*_0_ > 0) resp. suitable (*r*_0_ > 0) among species with different persistence ability.

On small spatial scales, niche-distribution mismatches can arise from source-sink effects and from time-delayed extinction (4-6). To assess the match between demographic suitability and a species’ occurrence within its geographic range, we regressed spatial variation in occupancy (on a 1’ grid, c. 1.55 × 1.85 km^2^) against the locally predicted *r*_0_. Occupancy generally increased with predicted *r*_0_ (Fig. 3b). But the strength of this relationship varied strongly among species (R^2^_N_: 0.01–0.55, mean = 0.18; Tab. S2) and significantly differed between species of different persistence ability, where variation in occupancy was better explained for nonsprouters than for resprouters (ANOVA, F_1,24_ = 4.74, p = 0.039). For resprouters, small-scale niche-distribution mismatches are greater mainly because populations more frequently occur in unsuitable sites (Fig. 3c). This can be explained by populations of more persistent species being less vulnerable to adverse conditions in both temporally fluctuating (29) and directionally changing environments (30). Dispersal ability had no positive effect on the occupancy of unsuitable sites. Hence, we found no indication of source-sink effects at the spatial scale of our analysis.

## Conclusions

In summary, our comparative analysis of demography-based niches indicates that key life history traits shape the geographic distributions of plant species by affecting not only niche sizes but also niche-distribution mismatches. Specifically, mismatches between niches and geographic distributions arise because poorly dispersed species are absent from suitable sites beyond their range limits and because species with high persistence ability are present in sites that are unsuitable under current, average environmental conditions. Importantly, this identifies poorly dispersing and highly persistent species as cases where static, correlative species distribution models are more likely to fail. For such species, range forecasts require dynamic species distribution models that incorporate demographic niche estimates (7, 8, 11). From a theoretical perspective, the quantification of spatial variation in species’ intrinsic population growth rates is also the first step towards understanding effects of biotic interactions on range dynamics, large-scale species coexistence (31, 32), and niche evolution (4). A demographic quantification of ecological niches, particularly in well-studied model systems, thus holds great promise for better integrating ecological theory and empirical biogeography. This is urgently needed to advance our predictive understanding of biodiversity dynamics under environmental change (10, 12).

## Materials and Methods

### Study species and demographic data

We studied 26 species of the Proteaceae family, specifically of the genera *Protea* (16 species) and *Leucadendron* (10 species), that are endemic to the Cape Floristic Region (17). These species were chosen to represent variation in geographic distributions as well as variation in dispersal and resprouting ability. For each species we obtained data on between-population variation in key demographic rates across the entire life cycle, namely the total fecundity of adult plants since the last fire (size of individual canopy seed banks), per capita post-fire seedling recruitment (ratio between post-fire recruits and pre-fire adults) and adult fire survival. The latter two rates were measured on recently burned sites (<3 years after fire), where burned pre-fire adults were still identifiable (33, 34). Study sites for demographic sampling were selected to cover major environmental gradients across the global geographic distribution of each study species. The final data set comprised 3,617 population-level records from an average of 99 (median = 85) study sites per species (Tab. S1). For details on the demographic data collection see ref. 19.

### Study region and environmental variables

Our study area was defined on a regular grid with a spatial resolution of 1’ × 1’ (c. 1.55 km × 1.85 km) and included all grid cells of the Cape Floristic Region in which > 5% of the area is covered by Fynbos vegetation (35). Climatic and edaphic variables that are expected to be main determinants of the performance and survival of serotinous Proteaceae were extracted from the South African Atlas of Climatology and Agrohydrology (36). We included *January maximum daily temperature* (T_max_), *July minimum daily temperature* (T_min_) and a *January aridity index* (AI) calculated as the ratio between the mean values of precipitation (P) and temperature (T): AI = P/(T + 10°C) (37). Climatic variables are averages over the years 1950–2000. As an edaphic variable we used a *soil fertility* index that combines soil texture and base status and ranges from 0 to 10 (36). Information on the fire return interval was obtained from both observational records and model predictions. For the demographic sampling sites, information on the fire history (time since the last fire and length of the previous fire interval) was inferred from a combination of measured plant ages, historical records and MODIS satellite observations (19, 38-40). For predictions of population growth rates across the study region (see below), we used probability distributions of fire return intervals predicted from a climate-driven model of post-fire ecosystem recovery (22).

### Demographic response model

We used a hierarchical Bayesian modelling approach for estimating the species-specific responses of key demographic rates (fecundity, per-seed establishment and adult fire survival) to environmental covariates (Fig. S3). The model considers effects of climatic and edaphic conditions, variable fire return intervals and intraspecific density dependence at both the adult and the seedling stage. Below, we describe the submodels for variation in each demographic rate.

#### Fecundity

The recorded size of the canopy seed bank (*Seed.count*_*i,j*_) of plant *j* in population *i* is described by an overdispersed Poisson distribution

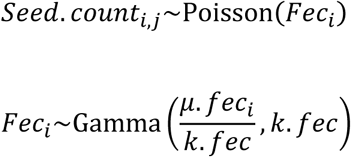

where the expected value of mean fecundity *μ.fec*_*i*_ is determined by limiting effects of post-fire stand age (*Age*), environmental covariates (**X**) and population density(*D*):

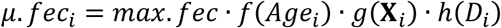

Effects of stand age on fecundity arise from the time of maturation until the first flowering and cone production, increasing accumulation of standing cones on growing plants, cone loss and possibly senescence of aged individuals:

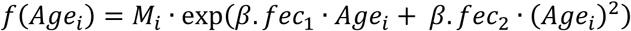

where *M*_i_ is a binary random variable (0, 1) indicating maturity. The probability of population-level maturity is calculated from a Weibull distribution for the age (*t.mat*) of first cone production:

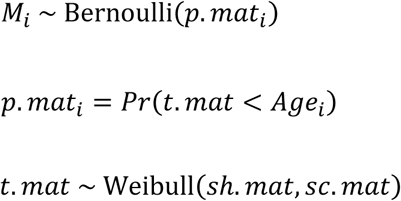

The species-specific time to reproductive maturity (*t.mat*) was constrained to be at least three years for nonsprouters (17). The effects of the environmental covariates *k* = 1…*K* are described by Gaussian demographic response functions:

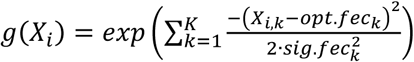

where *opt.fec*_*k*_ denotes the optimal conditions and *sig.fec*_*k*_ measures the width of the response curve. Effects of population density *D*_*i*_ on fecundity are described as (41):

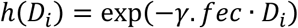

#### Establishment

The establishment of new recruits from seeds is modelled as a binomial process where the number of recruits (#*Recruits*_*i*_) in population *i* depends on the total number of available seeds (#*Seeds*_*i*_) in the canopy seed bank at the time of the last fire and the per-seed establishment rate π.*est*_*i*_:

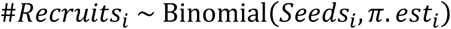

Since #*Seeds*_*i*_ is unknown for recently burned sites where recruitment was recorded, it is modelled as a latent state variable

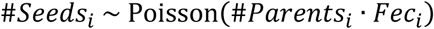

where #*Parents*_*i*_ denotes the number of pre-fire seed sources (only females for dioecious *Leucadendron* species) and *Fec*_*i*_ depends on environmental covariates (**X**_*i*_), the post-fire stand age (*Age*_*i*_) and the adult population density (*D*_*i*_) at the time of the previous fire as described in the fecundity submodel. Establishment rate π.*est*_*i*_ is affected by environmental covariates (**X**_*i*_) and by the densities of seeds *SD*_*i*_ = #*Seeds*_*i*_/*Area*_*i*_ and fire-surviving adults *AD*_*i*_ = #*Adults*_*i*_/*Area*_*i*_.

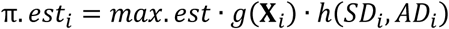

As for fecundity, the effects of different environmental covariates *k* = 1…*K* are described by Gaussian demographic response functions:

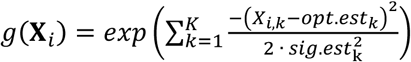

Density effects on establishment result from the density of seeds (*SD*_*i*_) as well of from the density of fire-surviving adults (*AD*_*i*_), with different strengths (γ.*est.SD* resp. γ.*est.AD*) for each of these density effects:

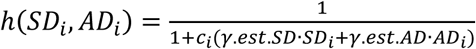

Density-dependent mortality of recruits (self-thinning) is a continuous process, which for Fynbos Proteaceae generally occurs within the first three years after a fire (42). This is described by weighting the density effects with a factor *c*_i_ that depends on the post-fire stand age (*pf.Age*_*i*_) at the time of sampling:

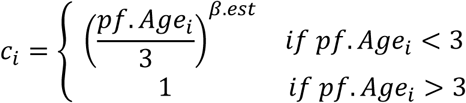

Thereby the model accounts for the fact that more seedlings can be observed if a site is surveyed just shortly after germination (19).

#### Survival

Adult fire survival is modelled as a binomial process for the proportion of survivors among all pre-fire adults (#*All.Adults*_*i*_):

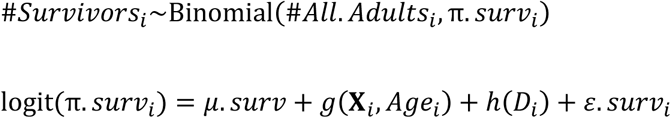

Similar as for fecundity, the effects of different environmental covariates *k* = 1…*K* and of post-fire stand age (*Age*_*i*_) are described by Gaussian response functions and effects of population density *D*_*i*_ by a negative-exponential function:

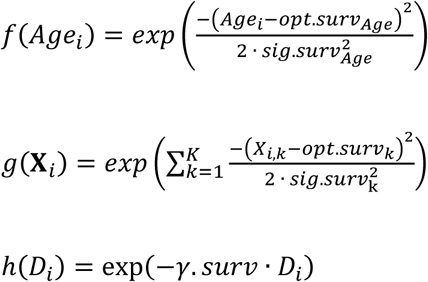

Since adult fire survival rates of nonsprouters are generally low with little intraspecific variation (19), we modelled them as species-specific constants and considered effects of covariates only for the survival rates of resprouters.

#### Bayesian parameter estimation

Parameters of the model were estimated independently for each study species. All environmental variables were scaled and centred and the *aridity index* and *soil fertility index* were additionally log-transformed before the analyses. The hierarchical model was formulated in a Bayesian framework and samples from the parameter posterior distribution were generated with Markov chain Monte Carlo (MCMC) methods in the software JAGS (43) (see *Supplementary Information* for model code). An overview of parameter prior distributions is given in Tab. S5. In three independent MCMC chains, posteriors were sampled from 100,000 iterations after a burn-in period of 500,000 iterations. Convergence of the MCMC sampler was checked by the multivariate scale reduction factor being smaller than 1.1 (44). For all further analyses, the posterior samples were regularly thinned to a sample size of 1,000 for each chain, resp. 3,000 samples in total.

#### Model evaluation

For each species we assessed the model fit separately for each observed demographic variable (fecundity, recruit:parent ratio, adult fire survival) by calculating Nagelkerke’s general 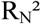 (45) relative to null models in which demographic rates (*π*.*est, π*.*surv, μ.fec*) are species-specific constants. The explained variance in each demographic variable for each study species is shown in Tab. S2.

### Prediction of niche sizes and geographic variation in *r*_0_

The demographic response model predicts fire survival rate *π*.*surv*, fecundity *μ.fec* and establishment rate *π*.*est* as functions of environmental covariates **X**, fire interval *T* and population density at different stages. Based on these demographic rates the expected population size *N* after a fire interval of length *T* can be calculated as the sum of fire survivors and new recruits:

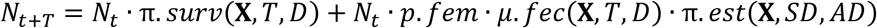

For the dioecious *Leucadendron* species the parameter *p.fem* specifies the proportion of female individuals in a population and accounts for the fact that fecundity rate *μ.fec* was defined per female. The niche of a species is defined as the set of environmental conditions for which the intrinsic growth rate of small populations (*r*_0_) is positive. To calculate *r*_0_ we set all density variables to zero and first calculated the rate of change in population size per fire interval

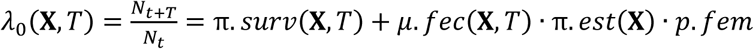

The intrinsic growth rate *r*_0_ was then calculated on an annual basis as

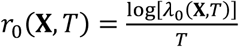

To quantify species’ niches, *r*_0_ was predicted on a 5-dimensional grid spanned by niche axes according to the four climatic-edaphic covariates (T_max_, T_min_, AI, soil fertility) and the fire return interval (log-transformed). Since the demographic data analysed in the demographic response models were measured in natural communities and thus incorporate effects of interspecific biotic interactions, the predicted *r*_0_ represents the post-interactive (or ‘realised’) niche (2). For commensurability of the different niche axes, these were confined and scaled to the respective range of values that occur throughout the Fynbos biome for each variable (so that each axis ranges from zero to one) and each niche axes was regularly sampled with a resolution of 0.01 (10^10^ grid points). The niche size was then quantified separately for fire return interval (‘disturbance niche size’) and for the four climatic-edaphic variables (‘environmental niche size’). The disturbance niche size was determined as the range of fire return intervals for which a positive *r*_0_ is predicted when the climatic-edaphic variables are set to their optimal values. Likewise, the environmental niche size was determined as the hypervolume in the 4-dimensional climatic-edaphic sub-space for which the predicted *r*_0_ is positive when setting the fire return interval to its species-specific optimum.

To geographically project *r*_0_ across the study region, *r*_0_ was not predicted for a single fixed fire return interval but integrated as a weighted geometric mean over the site-specific probability distribution of fire return intervals^20^. In all cases, values of *r*_0_ were predicted from each posterior sample of the demographic response model, yielding full posterior distributions for each predicted value.

### Species distribution data and geographic ranges

The Protea Atlas Project was an extensive citizen science project that used a standardized protocol to collected complete species lists from Proteaceae communities across the Cape Floristic Region (17). For our study region, the Protea Atlas data base contains 54,642 sampling locations with a total of 126,690 recorded presences of the 26 study species (Tab. S1). We aggregated these occurrence data to the proportion of presence records (occupancy) among the sampled communities in each 1’ × 1’ grid cell. As an overall assessment of how well species occurrence is predicted by demographic suitability, we calculated the area under the receiver operating characteristic curve (AUC) (46) for a binary classification of grid cell presence-absence by the predicted *r*_0_. We furthermore described the large-scale geographic range of each species by an alpha-convex hull over all presence records, using the *alphahull* package in R with parameter α = 0.5 (47, 48). The size of each species’ geographic range was calculated as the overlap of this alpha-convex hull with the Fynbos biome (study region). The degree of range filling was calculated as the proportion of potentially suitable area (cells with predicted *r*_0_>0) that lies within the geographic range. This calculation was conducted for each posterior sample of predicted *r*_0_ in order to account for uncertainty in estimated range filling.

### Statistical analyses of relationships between life history traits, niche characteristics and species geographic ranges

For the two measures of niche size (disturbance niche size and environmental niche size) as well as for species’ geographic range size and range filling, we performed separate regression analyses to test for effects of key life-history traits. The traits considered as explanatory variables were persistence ability (0 = nonsprouter, 1 = resprouter) and long-distance dispersal ability. The species-specific relative long-distance dispersal ability was derived from a trait-based mechanistic model of primary and secondary wind-dispersal (23) and measured as the number of neighbouring cells on a 1’ × 1’ rectangular grid that can be reached by dispersal from a source cell with a probability of at least 10^-4^. This dispersal measure was then log-transformed and scaled.

In each regression model we accounted for both measurements errors (posterior variance of the respective response variable) and phylogenetic dependence (49). Hence, the likelihood of the estimated species-specific posterior means (**Y.mean**) of the response variable depends on the vector of its true (but unknown) values **Y** and the respective posterior variances (**Y.var**):

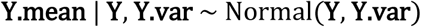

For the true **Y** we then formulated a multivariate normal model

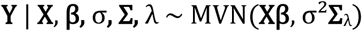

where **Xβ** is the linear predictor of the regression and **Σ**_λ_ is an adjusted variance-covariance matrix to account for phylogenetic dependencies. We first calculated the variance-covariance matrix **Σ** under a Brownian motion model using R package ape (50). A molecular phylogeny of our study species (Fig. S4) was obtained by pruning a phylogenetic tree of 291 Proteaceae species. This phylogenetic tree was constructed from a supermatrix combining molecular markers for *Leucadendron, Protea* and related species of the Proteaceae family (51-53). As quantitative measure of the degree of phylogenetic dependence the model furthermore includes Pagel’s λ (ranging from zero to one) (54) and the adjusted **Σ**_λ_ is calculated as **Σ**_λ_ **=** λ**Σ** + (1 – λ)**I**, where **I** is the identity matrix.

Bayesian parameter estimations were performed in JAGS using largely uninformative prior distributions (see *Supplementary Information* for model code). For each model we ran three independent MCMC chains with 200,000 iterations, the first half of which was discarded as burn-in and convergence was checked by the multivariate scale reduction factor being smaller than 1.1. For each response variable, we estimated a full model that included effects of persistence ability, dispersal ability and their interaction. All simplified models nested in the full model were then compared by the deviance information criteria (DIC) (55) and we report parameter estimates for the DIC-minimal models (Tab. S3).

### Statistical analysis of the relationship between occupancy and demographic suitability

We used a binomial non-linear regression model to analyse the relationship between each species’ occupancy in the 1’ × 1’ grid cells within its geographic range and the respectively predicted intrinsic population growth rate *r*_0_. The model describes the number of presence records (*y*_s,i_) for species *s* in the grid cell *i* as

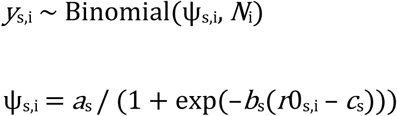

where *N*_i_ is the number of Protea Atlas sampling locations within the grid cell and *a*_s_, *b*_s_, *c*_s_ are species-specific regression parameters. Bayesian parameter estimation was performed in JAGS using largely uninformative prior distributions (see *Supplementary Information* for model code). We ran three independent MCMC chains with 20,000 iterations, the first half of which was discarded as burn-in and convergence was checked by the multivariate scale reduction factor being smaller than 1.1.

### Data and code availability

Phylogenetic data is available from the TreeBASE repository (http://purl.org/phylo/treebase/phylows/study/TB2:S23259). Demographic data were partly used under license from CapeNature for the current study, and so are not publicly available. These data are however available from the authors upon reasonable request and with permission of CapeNature. Documented JAGS code for the statistical analyses is provided in Supplementary Information. Additionally used R code and data generated during the analyses is available from the corresponding author upon request.

## Acknowledgements

This work was primarily funded by the German Research Foundation (DFG, grant SCHU 2259/5-1). M.T. acknowledges additional funding from Stellenbosch University and the DST-NRF’s Professional Development Programme (South Africa). We are grateful to CapeNature, SANParks and R.M. Cowling for access to demographic data and to B. Olivier, all field assistants, reserve managers and private landholders who supported our field work. Data were collected under the CapeNature permit AAA0028-AAA005-00213, Eastern Cape Parks permit CRO91/12CR and SANParks permit (Agulhas National Park). Many thanks to C. Buchmann, H. Cooksley, C.S. Sheppard and J. Walter for commenting on earlier drafts. We also thank Jake Alexander for helpful discussion and comments on a previous version of the manuscript.

## Author Contributions

J.P. and F.M.S. conceived and designed the study. M.T. collected the data with contributions from W.J.B., T.K., H.N., AL.S.-V. and F.M.S.. J.T. provided phylogenetic analyses. J.P. analysed the data and wrote the first draft. All co-authors contributed to the writing of the final manuscript.

## References

1. Hutchinson GE (1957) Concluding remarks. Cold Spring Harbor Symposia on Quantitative Biology 22:415–427.

2. Hutchinson GE (1978) An Introduction to Population Ecology (Yale University Press).

3. Maguire B (1973) Niche response structure and analytical potentials of its relationship to habitat. Am Nat 107:213–246.

4. Holt RD (2009) Bringing the Hutchinsonian niche into the 21st century: ecological and evolutionary perspectives. Proc Natl Acad Sci USA 106:19659–19665.

5. Schurr FM, et al. (2012) How to understand species’ niches and range dynamics: a demographic research agenda for biogeography. J Biogeogr 39:2146–2162.

6. Pulliam HR (2000) On the relationship between niche and distribution. Ecol Lett 3:349–361.

7. Pagel J, Schurr FM (2012) Forecasting species ranges by statistical estimation of ecological niches and spatial population dynamics. Glob Ecol Biogeogr 21:293–304.

8. Zurell D, et al. (2016) Benchmarking novel approaches for modelling species range dynamics. Glob Change Biol 22:2651–2664.

9. Settele J, et al. (2015) Terrestrial and inland water systems. Climate Change 2014 Impacts, Adaptation and Vulnerability: Part A: Global and Sectoral Aspects (Cambridge University Press), pp 271–359.

10. Ehrlén J, Morris WF (2015) Predicting changes in the distribution and abundance of species under environmental change. Ecol Lett 18:303–314.

11. Evans MEK, Merow C, Record S, McMahon SM, Enquist BJ (2016) Towards process-based range modeling of many species. Trends Ecol Evol 31:860–871.

12. Urban MC, et al. (2016) Improving the forecast for biodiversity under climate change. Science 353:aad8466.

13. Diez JM, Giladi I, Warren R, Pulliam HR (2014) Probabilistic and spatially variable niches inferred from demography. J Ecol 102:544–554.

14. Merow C, et al. (2014) On using integral projection models to generate demographically driven predictions of species’ distributions: development and validation using sparse data. Ecography 37:1167–1183.

15. Pironon S, et al. (2018) The ‘Hutchinsonian niche’ as an assemblage of demographic niches: implications for species geographic ranges. Ecography 41:1103–1113.

16. Myers N, Mittermeier RA, Mittermeier CG, Fonseca GAB, Kent J (2000) Biodiversity hotspots for conservation priorities. Nature 403:853–859.

17. Rebelo AG (2001) Proteas: A Field Guide to the Proteas of Southern Africa (Fernwood Press, Vlaeberg, South Africa).

18. Bond WJ, Van Wilgen BW (1996) Fire and Plants. Population and Community Biology Series 14 (Chapman & Hall).

19. Treurnicht M, et al. (2016) Environmental drivers of demographic variation across the global geographical range of 26 plant species. J Ecol 104:331–342.

20. Doak DF, Morris WF (2010) Demographic compensation and tipping points in climate-induced range shifts. Nature 467:959.

21. Villellas J, Doak DF, García MB, Morris WF (2015) Demographic compensation among populations: what is it, how does it arise and what are its implications? Ecol Lett 18:1139–1152.

22. Wilson AM, Latimer AM, Silander JA (2015) Climatic controls on ecosystem resilience: postfire regeneration in the Cape Floristic Region of South Africa. Proc Natl Acad Sci USA 112:9058–9063.

23. Schurr FM, et al. (2007) Colonization and persistence ability explain the extent to which plant species fill their potential range. Glob Ecol Biogeogr 16:449–459.

24. Thompson K, Gaston KJ, Band SR (1999) Range size, dispersal and niche breadth in the herbaceous flora of central England. J Ecol 87:150–155.

25. Lester SE, Ruttenberg BI, Gaines SD, Kinlan BP (2007) The relationship between dispersal ability and geographic range size. Ecol Lett 10:745–758.

26. Gaston KJ (2003) The Structure and Dynamics of Geographic Ranges (Oxford University Press).

27. Hargreaves AL, Karen ES, Eckert CG (2013) Are species’ range limits simply niche limits writ large? A review of transplant experiments beyond the range. Am Nat 183: 157–173.

28. Latimer AM, Silander JA, Rebelo AG, Midgley GF (2009) Experimental biogeography: the role of environmental gradients in high geographic diversity in Cape Proteaceae. Oecologia 160:151–162.

29. Higgins SI, Pickett ST, Bond WJ (2000) Predicting extinction risks for plants: environmental stochasticity can save declining populations. Trends Ecol Evol 15:516–520.

30. Enright NJ, Fontaine JB, Lamont BB, Miller BP, Westcott VC (2014) Resistance and resilience to changing climate and fire regime depend on plant functional traits. J Ecol 102:1572–1581.

31. Godsoe W, Jankowski J, Holt RD, Gravel D (2017) Integrating biogeography with contemporary niche theory. Trends Ecol Evol 32:488–499.

32. Usinowicz J, Levine JM (2018) Species persistence under climate change: a geographical scale coexistence problem. Ecol Lett 21:1589–1603.

33. Bond WJ, Vlok, J, Viviers M (1984) Variation in seedling recruitment of Cape Proteaceae after fire. J Ecol 72:209–221.

34. Bond WJ, Maze K, Desmet P (1995) Fire life histories and the seeds of chaos. Écoscience 2:252–260.

35. South African National Biodiversity Institute (2012) Vegetation Map of South Africa, Lesotho and Swaziland 2012 (http://bgis.sanbi.org/SpatialDataset/Detail/18). [vector geospatial dataset, downloaded on 11 August 2017]

36. Schulze RE (2007) South African Atlas of Climatology and Agrohydrology, Technical Report 1489/1/06 (Water Research Commission, Pretoria, South Africa).

37. De Martonne E (1926) Aréisme et indice artidite. C R Acad Sci 182:1395–1398.

38. De Klerk H (2008) A pragmatic assessment of the usefulness of the MODIS (Terra and Aqua) 1-km active fire (MOD14A2 and MYD14A2) products for mapping fires in the fynbos biome. Int J Wildland Fire 17:166–178.

39. Kraaij T, Cowling RM, Van Wilgen BW, Schutte-Vlok A (2013) Historical fire-regimes in a poorly understood, fire-prone ecosystem: eastern coastal fynbos. Int J Wildland Fire 22:277–287.

40. Roy DP, Boschetti L, Justice CO, Ju J (2008) The collection 5 MODIS burned area product – Global evaluation by comparison with the MODIS active fire product. Remote Sens Environ 112:3690–3707.

41. Nottebrock H, et al. (2017) Coexistence of plant species in a biodiversity hotspot is stabilized by competition but not by seed predation. Oikos 126:276–284.

42. Manders P, Smith R (1992) Effects of artificially established depth to water table gradients and soil type on the growth of Cape fynbos and forest plants. S Afr J Bot 58:195–201.

43. Plummer M (2003) JAGS: A program for analysis of Bayesian graphical models using Gibbs sampling. Proceedings of the 3rd International Workshop on Distributed Statistical Computing

44. Gelman A, Rubin DB (1992) Inference from iterative simulation using multiple sequences. Stat Sci 7:457–472.

45. Nagelkerke NJ (1991) A note on a general definition of the coefficient of determination. Biometrika 78:691–692.

46. Hanley JA, McNeil BJ (1982) The meaning and use of the area under a receiver operating characteristic (ROC) curve. Radiology 143:29–36.

47. Burgman MA, Fox JC (2003) Bias in species range estimates from minimum convex polygons: implications for conservation and options for improved planning. Anim Conserv 6:19–28.

48. Pateiro-Lopez B, Rodriguez-Casal A (2010) Generalizing the convex hull of a sample: the R package alphahull. J Stat Softw 34:1–28.

49. de Villemereuil P, Wells JA, Edwards RD, Blomberg SP (2012) Bayesian models for comparative analysis integrating phylogenetic uncertainty. BMC Evol Biol 12:102.

50. Paradis E, Claude J, Strimmer K (2004) APE: analyses of phylogenetics and evolution in R language. Bioinformatics 20:289–290.

51. Tonnabel J, et al. (2014) Convergent and correlated evolution of major life-history traits in the angiosperm genus *Leucadendron* (Proteaceae). Evolution 68:2775–2792.

52. Valente LM, et al. (2010) Diversification of the African genus Protea (Proteaceae) in the Cape biodiversity hotspot and beyond: equal rates in different biomes. Evolution 64:745–760.

53. Sauquet H, et al. (2009) Contrasted patterns of hyperdiversification in Mediterranean hotspots. Proc Natl Acad Sci USA. 106:221–225.

54. Pagel M (1999) Inferring the historical patterns of biological evolution. Nature 401:877–884.

55. Gelman A, Hwang J, Vehtari A (2014) Understanding predictive information criteria for Bayesian models. Stat Comput 24:997–1016.

