## Supplementary Information for "The niche is not the range: Dispersal and persistence shape mismatches between ecological niches and geographic distributions of plants"

In this supplementary information we provide the JAGS model codes that were used for the implementation of Bayesian parameter estimation at different steps in our analysis:

1) Demographic response model
2) Statistical analyses of relationships between life history traits, niche characteristics and species geographic ranges
3) Statistical analysis of the relationships between occupancy and demographic suitability

For each model we present the code together with an overview of all included data variables, parameters and parameter prior distributions.

### 1) Demographic response model

Parameters of the demographic response model were estimated separately for each study species. An overview of the hierarchical model structure is shown in Fig. S3.

**Data variables for the demographic response model**

| | Variable name [dimensions] | Description | Text symbol |
|---|---|---|---|
| Fecundity data | `n.Fec` | number of populations with fecundity data | |
| | `Fec.FEC[n.Fec]` | total size of the canopy seedbank the of sampled individuals | |
| | `No_Plants.FEC[n.Fec]` | number of sampled individuals | |
| | `log_AI.FEC[n.Fec]` | log-transformed January aridity index | *AI* |
| | `min_temp_jul.FEC[n.Fec]` | July minimum daily temperature | $T_{min}$ |
| | `max_temp_jan.FEC[n.Fec]` | January maximum daily temperature | $T_{max}$ |
| | `log_soil_fert.FEC[n.Fec]` | log-transformed soil fertility index | log(*soil fertility)* |
| | `Age.FEC[n.Fec]` | stand age | *Age* |
| | `SP.dens.FEC[n.Fec]` | population density | *D* |
| Recruitment data | `n.SPR` | number of populations with recruitment data | |
| | `Seedlings.SPR[n.SPR]` | number of recruits | #*Recruits* |
| | `Parents.SPR[n.SPR]` | number of pre-fire parents | #*Parents* |
| | `Alive.SPR[n.SPR]` | number of post-fire (fire-surviving) adults | #*Adults* |
| | `area.SPR[n.SPR]` | sampled area | *Area* |
| | `log_AI.SPR[n.SPR]` | log-transformed January aridity index | *AI* |
| | `min_temp_jul.SPR[n.SPR]` | July minimum daily temperature | $T_{min}$ |
| | `max_temp_jan.SPR[n.SPR]` | January maximum daily temperature | $T_{max}$ |
| | `log_soil_fert.SPR[n.SPR]` | log-transformed soil fertility index | log(*soil fertility)* |
| | `log_PF_Age.SPR[n.SPR]` | log-transformed, normalized post-fire stand age | min[log(*pf.Age*/3), 0] |
| | `Age.SPR[n.SPR]` | length of the previous fire interval, i.e. pre-fire stand age | *Age* |
| | `SP.dens.SPR[n.SPR]` | pre-fire population density | *D* |
| Survival data | `n.Surv` | number of populations with survival data | |
| | `Alive.SURV[n.Surv]` | number of fire-surviving adults | #*Survivors* |
| | `All_Adults.SURV[n.Surv]` | total number of pre-fire adults | #*All.Adults* |
| | `log_AI.SURV[n.Surv]` | log-transformed January aridity index | *AI* |
| | `min_temp_jul.SURV[n.Surv]` | July minimum daily temperature | $T_{min}$ |
| | `max_temp_jan.SURV[n.Surv]` | January maximum daily temperature | $T_{max}$ |
| | `log_soil_fert. SURV[n.Surv]` | log-transformed soil fertility index | log(*soil fertility)* |
| | `Age.SURV[n.Surv]` | length of the previous fire interval, i.e. pre-fire stand age | *Age* |
| | `SP.dens.SURV[n.Surv]` | pre-fire population density | *D* |

In this table 'Variable name' refers to a variable in the JAGS code (see below) and 'Text symbol' refers to the corresponding notation in the Methods section, if applicable. Note that in the fecundity submodel the measurements of individual-level sizes canopy seed banks ($Seed.count_{i,j}$) were aggregated to the sum (`Fec.FEC`) over all sampled individuals per population (`No_Plants.FEC`) for numerical efficiency. Environmental covariates are organized as separate variables for each submodel, although different demographic data types were often collected for the same population. For the sample sizes (`n.Fec`, `n.SPR`, `n.Surv`) for each species see Tab. S1.

**Parameters and prior distributions for the demographic response model**

| | Parameter name | Variable name | Description | Prior distribution | Prior parameters |
|---|---|---|---|---|---|
| Fecundity | log(*max.fec*) | `fec.Intercept` | maximum fecundity (log) | Normal | μ = 0, $\sigma^2 = 10^4$ |
| | $opt.fec_k$ | `fec.opt.log_AI` `fec.opt.min_temp_jul` `fec.opt.max_temp_jan` `fec.opt.log_soil_fert` | environmental optima | Normal | μ = 0, $\sigma^2 = 10^4$ |
| | $1/sig.fec^2_k$ | `fec.sc.log_AI` `fec.sc.min_temp_jul` `fec.sc.max_temp_jan` `fec.sc.log_soil_fert` | environmental response strengths | Exponential | λ = 1 |
| | *β.fec* | `fec.Age` `fec.Age_2` | age effects | Double-Exponential | μ = 0, λ = 1 |
| | *sh.mat* | `zi.sh` | Weibull parameters for age of maturity | Gamma | α = 0.01, β = 0.01 |
| | *sc.mat* | `zi.sc` | | Gamma | α = 0.01, β = 0.01 |
| | *γ.fec* | `fec.SP.dens` | strength of density effects | Exponential | λ = 1 |
| | *k.fec* | `size.fec` | overdispersion parameter | Gamma | α = 0.01, β = 0.01 |
| Establishment | *max.est* | `recr.Intercept` | maximum establishment rate | Beta | a = 1, b = 1 |
| | $opt.est_k$ | `recr.opt.log_AI` `recr.opt.min_temp_jul` `recr.opt.max_temp_jan` `recr.opt.log_soil_fert` | environmental optima | Normal | μ = 0, $\sigma^2 = 10^4$ |
| | $1/sig.est^2_k$ | `recr.sc.log_AI` `recr.sc.min_temp_jul` `recr.sc.max_temp_jan` `recr.sc.log_soil_fert` | environmental response strengths | Exponential | λ = 1 |
| | *β.est* | `recr.log_PF_Age` | effect of time since fire | Exponential | λ = 1 |
| | *γ.est.SD* | `recr.seed.dens` | strength of density effects from seeds | Exponential | λ = 1 |
| | *γ.est.AD* | `recr.adult.dens` | strength of density effects from adults | Exponential | λ = 1 |
| Survival | *max.surv* | `surv.Intercept` | maximum survival rate | Beta | a = 1, b = 1 |
| | $opt.surv_k$ | `surv.opt.log_AI` `surv.opt.min_temp_jul` `surv.opt.max_temp_jan` `surv.opt.log_soil_fert` | environmental optima | Normal | μ = 0, $\sigma^2 = 10^4$ |
| | $1/sig.surv^2_k$ | `surv.sc.log_AI` `surv.sc.min_temp_jul` `surv.sc.max_temp_jan` `surv.sc.log_soil_fert` | environmental response strengths | Exponential | λ = 1 |
| | $opt.surv_{Age}$ | `surv.opt.Age` | age optimum | Normal | μ = 0, $\sigma^2 = 10^4$ |
| | $1/sig.surv_{Age}$ | `surv.sc.Age` | age response strengths | Exponential | λ = 1 |
| | *γ.surv* | `surv.SP.dens` | strength of density effects | Exponential | λ = 1 |

This is an adjusted version of Tab. S5. In this table ‘Parameter name’ refers to the notation in the Methods and ‘Variable name’ to the corresponding variable in the JAGS code (see below).

**JAGS code for the demographic response model**

```
model {
### Fecundity submodel
for (i in 1:n.Fec){
  # Negative-binomial model for the population-level seed count
  Fec.FEC[i] ~ dnegbin(prob.FEC[i], size.fec)
  prob.FEC[i] <- size.fec/(size.fec + mu.Fec[i] * No_Plants.FEC[i] * I.fec.FEC[i])
  I.fec.FEC[i] ~ dbern(zi.fec.FEC[i])

  # Age-dependent probability of maturity according to a Weibull model
  I.fec.FEC[i] ~ dbern(zi.fec.FEC[i])
  zi.fec.FEC[i] <- 1 - exp(-(Age.mat.FEC[i]/zi.sc)^zi.sh)

  # Calculation of effects of environment (Gaussian response curves), age and population density
  log(mu.Fec[i]) <- fec.Intercept - fec.sc.log_AI * pow(fec.opt.log_AI - log_AI.FEC[i], 2)
                   - fec.sc.min_temp_jul * pow(fec.opt.min_temp_jul - min_temp_jul.FEC[i], 2)
                   - fec.sc.max_temp_jan * pow(fec.opt.max_temp_jan - max_temp_jan.FEC[i], 2)
                   - fec.sc.log_soil_fert * pow(fec.opt.log_soil_fert - log_soil_fert.FEC[i], 2)
                   + fec.Age * Age.FEC[i] + fec.Age_2 * pow(Age.FEC[i],2)
                   - fec.SP.dens * SP.dens.FEC[i]
  }
```

```
### Establishment submodel
for (j in 1:n.SPR){
  # Negative-binomial model for number of observed recruits
  Seedlings.SPR[j] ~ dnegbin(prob.SPR[j], size.fec)
  prob.SPR[j] <- size.fec/(size.fec + Seeds[j] * p.recr[j])
  p.recr[j] <- recr.Intercept * mu.recr[j] * dens.fac[j]

  # Age-dependent density effects
  dens.fac[j] <- 1/(1 + c[j] * (recr.seed.dens * Seeds[j]/area.SPR[j] + recr.adult.dens *
                Alive.SPR[j]/area.SPR[j]))
  log(c[j]) <- recr.log_PF_Age * log_PF_Age.SPR[j]

  # Calculation of environmental effects (Gaussian response curves)
  log(mu.recr[j]) <-  - recr.sc.log_AI * pow(recr.opt.log_AI - log_AI.SPR[j], 2)
                      - recr.sc.min_temp_jul * pow(recr.opt.min_temp_jul - min_temp_jul.SPR[j], 2)
                      - recr.sc.max_temp_jan * pow(recr.opt.max_temp_jan - max_temp_jan.SPR[j], 2))
                      - recr.sc.log_soil_fert * pow(recr.opt.log_soil_fert - log_soil_fert.SPR[j], 2)


  # The expected number of seeds is predicted from the fecundity submodel (see above):
  Seeds[j] <- mu.Fec.SPR[j] * Parents.SPR[j] * I.fec.SPR[j]
  I.fec.SPR[j] ~ dbern(zi.fec.SPR[j])
  zi.fec.SPR[j] <- (1 -exp(-(Age.mat.SPR[j]/zi.sc)^zi.sh))
  log(mu.Fec.SPR[j]) <- fec.Intercept - fec.sc.log_AI * pow(fec.opt.log_AI - log_AI.SPR[j], 2)
                        - fec.sc.min_temp_jul * pow(fec.opt.min_temp_jul - min_temp_jul.SPR[j], 2)
                        - fec.sc.max_temp_jan * pow(fec.opt.max_temp_jan - max_temp_jan.SPR[j], 2)
                        - fec.sc.log_soil_fert * pow(fec.opt.log_soil_fert - log_soil_fert.SPR[j] , 2)
                        + fec. Age * Age.SPR[j] + fec. Age_2 * pow(Age.SPR[j], 2)
                        - fec.SP.dens * SP.dens.SPR[j]
  }

### Survival submodel
for (k in 1:n.Surv){
  # Binomial model for number of surviving adults
  Alive.SURV[k] ~ dbin(p.surv[k], All_Adults.SURV[k])
  p.surv[k] <- mu.surv[k] * surv.Intercept
  # Calculation of environmental effects from Gaussian response curves
  log(mu.surv[k]) <-  - surv.sc.log_AI * pow(surv.opt.log_AI - log_AI.SURV[k], 2)
                      - surv.sc.min_temp_jul * pow(surv.opt.min_temp_jul - min_temp_jul.SURV[k], 2)
                      - surv.sc.max_temp_jan * pow(surv.opt.max_temp_jan - max_temp_jan.SURV[k], 2)
                      - surv.sc.log_soil_fert * pow(surv.opt.log_soil_fert - log_soil_fert.SURV[k], 2
                      - surv.sc. Age * pow(surv.opt. Age - Age.SURV[k], 2)
                      - surv.SP.dens * SP.dens.SURV[j]
  }

### Prior distributions
fec.opt.log_AI ~ dnorm(0, 0.0001)
fec.opt.min_temp_jul ~ dnorm(0, 0.0001)
fec.opt.max_temp_jan ~ dnorm(0, 0.0001)
fec.opt.log_soil_fert ~ dnorm(0, 0.0001)
recr.opt.log_AI ~ dnorm(0, 0.0001)
recr.opt.min_temp_jul ~ dnorm(0, 0.0001)
recr.opt.max_temp_jan ~ dnorm(0, 0.0001)
recr.opt.log_soil_fert ~ dnorm(0, 0.0001)
surv.opt.log_AI ~ dnorm(0, 0.0001)
surv.opt.min_temp_jul ~ dnorm(0, 0.0001)
surv.opt.max_temp_jan ~ dnorm(0, 0.0001)
surv.opt.log_soil_fert ~ dnorm(0, 0.0001)
surv.opt.Age ~ dnorm(0, 0.0001)
fec.sc.log_AI ~ dexp(1)
fec.sc.min_temp_jul ~ dexp(1)
fec.sc.max_temp_jan ~ dexp(1)
fec.sc.log_soil_fert ~ dexp(1)
recr.sc.log_AI ~ dexp(1)
recr.sc.min_temp_jul ~ dexp(1)
recr.sc.max_temp_jan ~ dexp(1)
recr.sc.log_soil_fert ~ dexp(1)
surv.sc.log_AI ~ dexp(1)
surv.sc.min_temp_jul ~ dexp(1)
surv.sc.max_temp_jan ~ dexp(1)
surv.sc.log_soil_fert ~ dexp(1)
surv.sc.Age ~ dexp(1)
fec.Age ~ ddexp(0,1)
fec.Age_2 ~ ddexp(0,1) T(,0)
fec.SP.dens ~ dexp(1)
recr.seed.dens ~ dexp(1)
recr.adult.dens ~ dexp(1)
recr.log_PF_Age ~ dexp(1)
surv.SP.dens ~ dexp(1)
zi.sc ~ dgamma(0.01, 0.01)T(0.1,)
zi.sh ~ dgamma(0.01, 0.01)T(0.1,)
surv.Intercept ~ dbeta(1,1)
fec.Intercept ~ dnorm(0, 0.0001)
recr.Intercept ~ dbeta(1,1)
size.fec ~ dgamma(0.01,0.01)
}
```

#### 2) Statistical analyses of relationships between life history traits, niche characteristics and species geographic ranges

We used identical model structures for analysing effects of life history traits on different response variables (disturbance niche size, environmental niche size, potential range size, range filling, range size, see Tab. S3). In each case, we formulated a full model that included effects of persistence ability, dispersal ability and their interaction as well as all simplified models nested within this full model. Here we only document the full model.

**Data variables for the regression of niche and range characteristics against life history traits**

| Variable name [dimensions] | Description | Text symbol |
|---|---|---|
| `n.Spec` | number of species (26) | |
| `Y.mean[n.Spec]` | response variable (posterior means) | **Y.mean** |
| `Y.Var[n.Spec, n.Spec]` | uncertainty of the response variable (posterior variances as diagonal matrix) | **Y.var** |
| `Disp[n.Spec]` | species' dispersal ability (log-transformed and scaled) | |
| `Pers[n.Spec]` | species' persistence ability (0 = nonsprouter, 1 = resprouter) | |
| `A[n.Spec, n.Spec]` | unadjusted covariance matrix | $\mathbf{\Sigma}$ |
| `ID[n.Spec, n.Spec]` | identity matrix | $\mathbf{I}$ |

In this table 'Variable name' refers to a variable in the JAGS code (see below) and 'Text symbol' refers to the corresponding notation in the Methods section, if applicable. Note that for the response variable 'range size' we have no quantification of interspecific variation in precision (i.e. posterior variances) and thus `Y.Var` was set to zero in those analyses.

**Parameters for the regression of niche and range characteristics against life history traits**

| Parameter name | Variable name | Description | Prior distribution | Prior parameters |
|---|---|---|---|---|
| $\beta_0$ | `alpha` | intercept | Normal | $\mu = 0, \sigma^2 = 10^4$ |
| $\beta_{Disp}$ | `beta.D` | effect of dispersal ability | Normal | $\mu = 0, \sigma^2 = 10^4$ |
| $\beta_{Pers}$ | `beta.P` | effect of persistence ability | Normal | $\mu = 0, \sigma^2 = 10^4$ |
| $\beta_{Disp:Pers}$ | `beta.DP` | interaction effect | Normal | $\mu = 0, \sigma^2 = 10^4$ |
| $\lambda$ | `lambda` | Pagel's λ | Beta | a = 2, b = 2 |
| $\sigma^2$ | `sig2` | residual variance | Inv.-Gamma | $\alpha = 0.01, \beta = 0.01$ |

In this table 'Parameter name' refers to the notation in the Methods (and Tab. S3) and 'Variable name' to the corresponding variable in the JAGS code (see below).

**JAGS code for the regression of niche and range characteristics against life history traits**

```
model {
# Multivariate normal model
Y.mean[1:n.Spec] ~ dmnorm(mu[],TAU[,])

# Linear predictor for each species
for (i in 1:n.Spec) {
  mu[i] <- alpha + beta.D*Disp[i] + beta.P*Pers[i] + beta.DP*Disp[i]*Pers[i]
  }

# Calculation of the adjusted, combined covariance matrix
Mlam <- sig2*(lambda*A[,] + (1-lambda)*ID) + Y.Var[,]
TAU <- inverse(Mlam)

### Prior distributions
alpha ~ dnorm(0, 1.0E-04)
beta.D ~ dnorm(0, 1.0E-04)
beta.P ~ dnorm(0, 1.0E-04)
beta.DP ~ dnorm(0, 1.0E-04)
lambda ~ dbeta(2,2)
tau ~ dgamma(0.01,0.01)
sig2 <- 1/tau
}
```

#### 3) Statistical analysis of the relationship between occupancy and demographic suitability

The nonlinear regression of grid-cell occupancy against demographic suitability (predicted $r_0$) is performed jointly for all species, while estimating species-specific values for the regression parameters ($a_s$, $b_s$, $c_s$). In order to allow predictions of the average relationships between occupancy and demographic suitability for different persistence abilities (Fig. 3b), the model includes separate hyperparameters for the mean of each regression parameter among nonsprouter resp. resprouter species.

**Data variables for the regression of occupancy against demographic suitability**

| Variable name [dimensions] | Description | Text symbol |
|---|---|---|
| `n.Dat` | number of data points (91288)<br>(species-grid cell combinations) | |
| `vis[n.Dat]` | number of presence-absence data (Protea Atlas records) in the grid cell | $N$ |
| `pres[n.Dat]` | number of recorded species presences in the grid cell | $y$ |
| `R[n.Dat]` | predicted demographic suitability | $r_0$ |
| `SP[n.Dat]` | numerical species index | $s$ |
| `n.Spec` | number of species (26) | |
| `Pers[n.Spec]` | species' persistence ability<br>(0 = nonsprouter, 1 = resprouter) | |

In this table 'Variable name' refers to a variable in the JAGS code (see below) and 'Text symbol' refers to the corresponding notation in the Methods section, if applicable.

**Parameters for the regression of occupancy against demographic suitability**

| Parameter name | Variable name | Description | Prior distribution | Prior parameters |
|---|---|---|---|---|
| $a$ | `a[n.Spec]` | species-specific regression parameter *a* | | |
| | `MU.a[2]` | mean of logit(*a*) for nonsprouters resp. sprouters | Normal | $\mu = 0, \sigma^2 = 10^4$ |
| | `sig2.a` | interspecific variation in logit(*a*) | Inv.-Gamma | $\alpha = 0.01, \beta = 0.01$ |
| $b$ | `b[n.Spec]` | species-specific regression parameter *b* | | |
| | `MU.b[2]` | mean of log(*b*) for nonsprouters resp. sprouters | Normal | $\mu = 0, \sigma^2 = 10^4$ |
| | `sig2.b` | interspecific variation in log(*b*) | Inv.-Gamma | $\alpha = 0.01, \beta = 0.01$ |
| $c$ | `c[n.Spec]` | species-specific regression parameter *c* | | |
| | `MU.c[2]` | mean of *c* for nonsprouters resp. sprouters | Normal | $\mu = 0, \sigma^2 = 10^4$ |
| | `sig2.c` | interspecific variation in *c* | Inv.-Gamma | $\alpha = 0.01, \beta = 0.01$ |

In this table 'Parameter name' refers to the notation in the Methods and 'Variable name' to the corresponding variable in the JAGS code (see below). Note that prior distributions are not specified for the species-specific regression parameters, but for the hyperparameters that describe interspecific variation in regression parameters.

**JAGS code for the regression of occupancy against demographic suitability**

```
model {
# Non-linear regression of occupancy against r0 for each species-site combination
for (i in 1: n.Dat) {
  pres[i] ~ dbin(psi[i],vis[i])
  psi[i] <- a[SP[i]]/(1 + exp(-b[SP[i]]*(R[i] - c[SP[i]])))
  }

# Interspecific variation in regression parameters a, b, c
for (sp in 1:n.Spec) {
  logit(a[sp]) <- logit.a[sp]
  logit.a[sp] ~ dnorm(mu.a[sp], tau.a)
  mu.a[sp] <- MU.a[Pers[sp] + 1]

  log(b[sp]) <- log.b[sp]
  log.b[sp] ~ dnorm(mu.b[sp], tau.b)
  mu.b[sp] <- MU.b[Pers[sp] + 1]

  c[sp] ~ dnorm(mu.c[sp], tau.c)
  mu.c[sp] <- MU.c[Pers[sp] + 1]
  }

### Prior distributions
MU.a[1] ~ dnorm(0,1.0E-04)
MU.a[2] ~ dnorm(0,1.0E-04)
tau.a  ~ dgamma(0.01,0.01)
sig2.a <- 1/tau.a
MU.b[1] ~ dnorm(0,1.0E-04)
MU.b[2] ~ dnorm(0,1.0E-04)
tau.b ~ dgamma(0.01,0.01)
sig2.b <- 1/tau.b
MU.c[1] ~ dnorm(0,1.0E-04)
MU.c[2] ~ dnorm(0,1.0E-04)
tau.c ~ dgamma(0.01,0.01)
sig2.c <- 1/tau.c
}
```

**Table S1. Overview of study species, samples sizes of demographic data and geographic distribution data**

| Species | Resprouting ability | No. sampled populations | Sample sizes of demographic data | | | No. presence records* | Geographic range size (km²)† |
|---|---|---|---|---|---|---|---|
| | | | Fecundity | Recruitment | Survival | | |
| *Leucadendron album* | nonsprouter | 48 | 41 | 22 | 15 | 672 | 8,720 |
| *Leucadendron coniferum* | nonsprouter | 68 | 45 | 23 | 0 | 793 | 3,075 |
| *Leucadendron eucalyptifolium* | nonsprouter | 70 | 19 | 51 | 0 | 5,248 | 26,157 |
| *Leucadendron laureolum* | nonsprouter | 75 | 68 | 28 | 23 | 3,453 | 8,658 |
| *Leucadendron modestum* | nonsprouter | 80 | 76 | 18 | 14 | 647 | 3,576 |
| *Leucadendron muirii* | nonsprouter | 80 | 70 | 16 | 6 | 599 | 2,956 |
| *Leucadendron rubrum* | nonsprouter | 136 | 69 | 77 | 17 | 4,634 | 36,220 |
| *Leucadendron salignum* | resprouter | 141 | 99 | 79 | 77 | 24,373 | 60,228 |
| *Leucadendron spissifolium* | resprouter | 90 | 80 | 37 | 36 | 4,676 | 35,653 |
| *Leucadendron xanthoconus* | nonsprouter | 85 | 61 | 38 | 16 | 7,170 | 5,999 |
| *Protea acaulos* | resprouter | 89 | 80 | 51 | 47 | 3,802 | 17,408 |
| *Protea amplexicaulis* | nonsprouter | 77 | 74 | 22 | 26 | 1,242 | 8,879 |
| *Protea compacta* | nonsprouter | 85 | 77 | 30 | 22 | 902 | 3,226 |
| *Protea cynaroides* | resprouter | 86 | 83 | 27 | 24 | 8,488 | 33,533 |
| *Protea eximia* | nonsprouter | 98 | 53 | 47 | 2 | 2,391 | 22,995 |
| *Protea laurifolia* | nonsprouter | 100 | 78 | 38 | 20 | 10,934 | 25,516 |
| *Protea longifolia* | nonsprouter | 84 | 78 | 34 | 28 | 1,635 | 5,234 |
| *Protea lorifolia* | nonsprouter | 142 | 54 | 91 | 4 | 5,259 | 24,904 |
| *Protea neriifolia* | nonsprouter | 150 | 68 | 95 | 16 | 6,382 | 30,129 |
| *Protea nitida* | resprouter | 83 | 76 | 30 | 30 | 9,943 | 42,426 |
| *Protea obtusifolia* | nonsprouter | 83 | 62 | 27 | 7 | 1,337 | 5,262 |
| *Protea punctata* | nonsprouter | 85 | 50 | 37 | 2 | 2,319 | 21,546 |
| *Protea repens* | nonsprouter | 292 | 104 | 224 | 42 | 15,291 | 59,077 |
| *Protea scabra* | resprouter | 92 | 85 | 65 | 65 | 2,254 | 5,970 |
| *Protea scolopendriifolia* | resprouter | 76 | 76 | 30 | 30 | 1,283 | 21,444 |
| *Protea susannae* | nonsprouter | 83 | 48 | 36 | 1 | 963 | 5,112 |

* Protea Atlas data base (Rebelo, 2001)

† Approximate geographic range size was calculated as overlap between an alpha-convex hull over the presence records of a species and the Fynbos biome (study region).

**Table S2. Explained intraspecific variation in demographic rates and occurrence**

| Species | Demographic rates ($R^2_N$) | | | Presence-absence (AUC) | Within-range occupancy ($R^2_N$) |
|---|---|---|---|---|---|
| | Fecundity | Recruitment | Survival* | | |
| *Leucadendron album* | 0.91 | 0.69 | - | 0.96 | 0.55 |
| *Leucadendron coniferum* | 0.38 | 0.46 | - | 0.95 | 0.30 |
| *Leucadendron eucalyptifolium* | 0.38 | 0.38 | - | 0.75 | 0.34 |
| *Leucadendron laureolum* | 0.29 | 0.45 | - | 0.87 | 0.01 |
| *Leucadendron modestum* | 0.43 | 0.84 | - | 0.93 | 0.46 |
| *Leucadendron muirii* | 0.34 | 0.36 | - | 0.96 | 0.22 |
| *Leucadendron rubrum* | 0.48 | 0.05 | - | 0.64 | 0.04 |
| *Leucadendron salignum* | 0.69 | 0.26 | 0.52 | 0.54 | 0.06 |
| *Leucadendron spissifolium* | 0.80 | 0.45 | 0.52 | 0.59 | 0.20 |
| *Leucadendron xanthoconus* | 0.56 | 0.44 | - | 0.92 | 0.27 |
| *Protea acaulos* | 0.53 | 0.32 | - | 0.69 | 0.01 |
| *Protea amplexicaulis* | 0.81 | 0.53 | - | 0.70 | 0.15 |
| *Protea compacta* | 0.59 | 0.66 | - | 0.93 | 0.05 |
| *Protea cynaroides* | 0.71 | 0.61 | 0.64 | 0.69 | 0.11 |
| *Protea eximia* | 0.67 | 0.48 | - | 0.70 | 0.12 |
| *Protea laurifolia* | 0.55 | 0.12 | - | 0.55 | 0.01 |
| *Protea longifolia* | 0.48 | 0.36 | - | 0.92 | 0.27 |
| *Protea lorifolia* | 0.62 | 0.30 | - | 0.77 | 0.07 |
| *Protea neriifolia* | 0.44 | 0.38 | - | 0.69 | 0.20 |
| *Protea nitida* | 0.70 | 0.23 | 0.41 | 0.57 | 0.17 |
| *Protea obtusifolia* | 0.51 | 0.50 | - | 0.95 | 0.26 |
| *Protea punctata* | 0.53 | 0.47 | - | 0.74 | 0.43 |
| *Protea repens* | 0.73 | 0.17 | - | 0.59 | 0.11 |
| *Protea scabra* | 0.63 | 0.51 | 0.81 | 0.53 | 0.02 |
| *Protea scolopendriifolia* | 0.69 | 0.50 | 0.83 | 0.73 | 0.01 |
| *Protea susannae* | 0.60 | 0.46 | - | 0.97 | 0.28 |

*Note that intraspecific variation in adult fire survival rates was modelled only for resprouter species but not for nonsprouter species (indicated by '-') that have very low fire survival rates with little intraspecific variation[19].

**Table S3. Effects of life history traits on the sizes of ecological niches and geographic ranges**

| Response variable | Model parameters | | | | | |
|---|---|---|---|---|---|---|
| | $\beta_0$ | $\beta_{Disp}$ | $\beta_{Pers}$ | $\beta_{Disp:Pers}$ | $\Lambda$ | $\sigma$ |
| Disturbance niche size (logit) | 0.55 ± 0.23 | - | **1.34 ± 0.41** | - | 0.82 ± 0.19 | 0.52 ± 0.11 |
| Environmental niche size (logit) | -1.42 ± 0.60 | **1.00 ± 0.40** | - | - | 0.30 ± 0.21 | 1.78 ± 0.42 |
| Potential range size (log) | 8.81 ± 0.35 | 0.30 ± 0.19 | -. | - | 0.40 ± 0.24 | 0.97 ± 0.20 |
| Range filling (logit) | 0.12 ± 0.49 | **0.58 ± 0.29** | -. | - | 0.45 ± 0.25 | 1.33 ± 0.27 |
| Range size (log) | 7.78 ± 0.39 | **0.46 ± 0.24** | **1.08 ± 0.54** | - | 0.23 ± 0.18 | 1.20 ± 0.21 |

Effects of life history traits on each of the different response variables were estimated by Bayesian normal linear regression analysis with model selection. Each full model included main effects of dispersal ability (*Disp*) and persistence ability (*Pers*; coded as 0 = nonsprouter, 1 = resprouter) as well as their interaction. All simplified models nested in the full model were compared by the deviance information criteria (DIC) and estimated parameters (posterior mean ± standard deviation) are shown for the respective DIC-minimal model ('-' indicates non-included terms). Significant regression coefficients (95% credibility interval does not overlap with zero) are in bold font. All models corrected for phylogenetic relatedness by estimating Pagel's Λ. σ denotes the residual standard deviation.

**Table S4. Model evaluation for populations established outside the species' geographic ranges**

| Species | No. populations | Proportion with $r_0 > 0$ |
|---|---|---|
| *Protea eximia* | 206 | 89% |
| *Protea compacta* | 162 | 35% |
| *Protea neriifolia* | 90 | 100% |
| *Protea laurifolia* | 48 | 100% |
| *Protea susannae* | 36 | 31% |
| *Leucadendron coniferum* | 25 | 20% |
| *Leucadendron laureolum* | 25 | 92% |
| *Protea lorifolia* | 21 | 81% |
| *Protea obtusifolia* | 13 | 62% |
| *Leucadendron eucalyptifolium* | 8 | 100% |
| *Protea longifolia* | 7 | 100% |
| *Leucadendron xanthoconus* | 6 | 67% |
| *Protea punctata* | 6 | 83% |
| *Protea cynaroides* | 4 | 75% |
| *Leucadendron rubrum* | 2 | 100% |
| *Leucadendron muirii* | 1 | 100% |
| *Protea amplexicaulis* | 1 | 100% |
| *Leucadendron album* | 0 | n.a. |
| *Leucadendron modestum* | 0 | n.a. |
| *Leucadendron salignum* | 0 | n.a. |
| *Leucadendron spissifolium* | 0 | n.a. |
| *Protea acaulos* | 0 | n.a. |
| *Protea nitida* | 0 | n.a. |
| *Protea repens* | 0 | n.a. |
| *Protea scabra* | 0 | n.a. |
| *Protea scolopendriifolia* | 0 | n.a. |

The table shows for each species the number of recorded populations that were established outside their respective geographic range[17]. We evaluated model predictions of the intrinsic population growth rate $r_0$ for the locations of these populations and report here for each species the proportion of populations for which a positive growth rate (demographic suitability) was predicted.

**Table S5. Prior distributions for parameters of the demographic response model**

| Model parameter | | Description | Prior distribution | Prior parameters |
|---|---|---|---|---|
| Fecundity | log(*max.fec*) | maximum fecundity (log) | Normal | $\mu = 0, \sigma^2 = 10^4$ |
| | $opt.fec_k$ | environmental optima | Normal | $\mu = 0, \sigma^2 = 10^4$ |
| | $1/sig.fec^2_k$ | environmental response strengths | Exponential | $\lambda = 1$ |
| | *β.fec* | age effects | Double-Exponential | $\mu = 0, \lambda = 1$ |
| | *sh.mat* | Weibull parameters for age of maturity | Gamma | $\alpha = 0.01, \beta = 0.01$ |
| | *sc.mat* | | Gamma | $\alpha = 0.01, \beta = 0.01$ |
| | *γ.fec* | strength of density effects | Exponential | $\lambda = 1$ |
| | *k.fec* | overdispersion parameter | Gamma | $\alpha = 0.01, \beta = 0.01$ |
| Establishment | *max.est* | maximum establishment rate | Beta | a = 1, b = 1 |
| | $opt.est_k$ | environmental optima | Normal | $\mu = 0, \sigma^2 = 10^4$ |
| | $1/sig.est^2_k$ | environmental response strengths | Exponential | $\lambda = 1$ |
| | *β.est* | effect of time since fire | Exponential | $\lambda = 1$ |
| | *γ.est.SD* | strength of density effects from seeds | Exponential | $\lambda = 1$ |
| | *γ.est.AD* | strength of density effects from adults | Exponential | $\lambda = 1$ |
| Survival | *max.surv* | maximum survival rate | Beta | a = 1, b = 1 |
| | $opt.surv_k$ | environmental optima | Normal | $\mu = 0, \sigma^2 = 10^4$ |
| | $1/sig.surv^2_k$ | environmental response strengths | Exponential | $\lambda = 1$ |
| | $opt.surv_{Age}$ | age optimum | Normal | $\mu = 0, \sigma^2 = 10^4$ |
| | $1/sig.surv_{Age}$ | age response strengths | Exponential | $\lambda = 1$ |
| | *γ.surv* | strength of density effects | Exponential | $\lambda = 1$ |

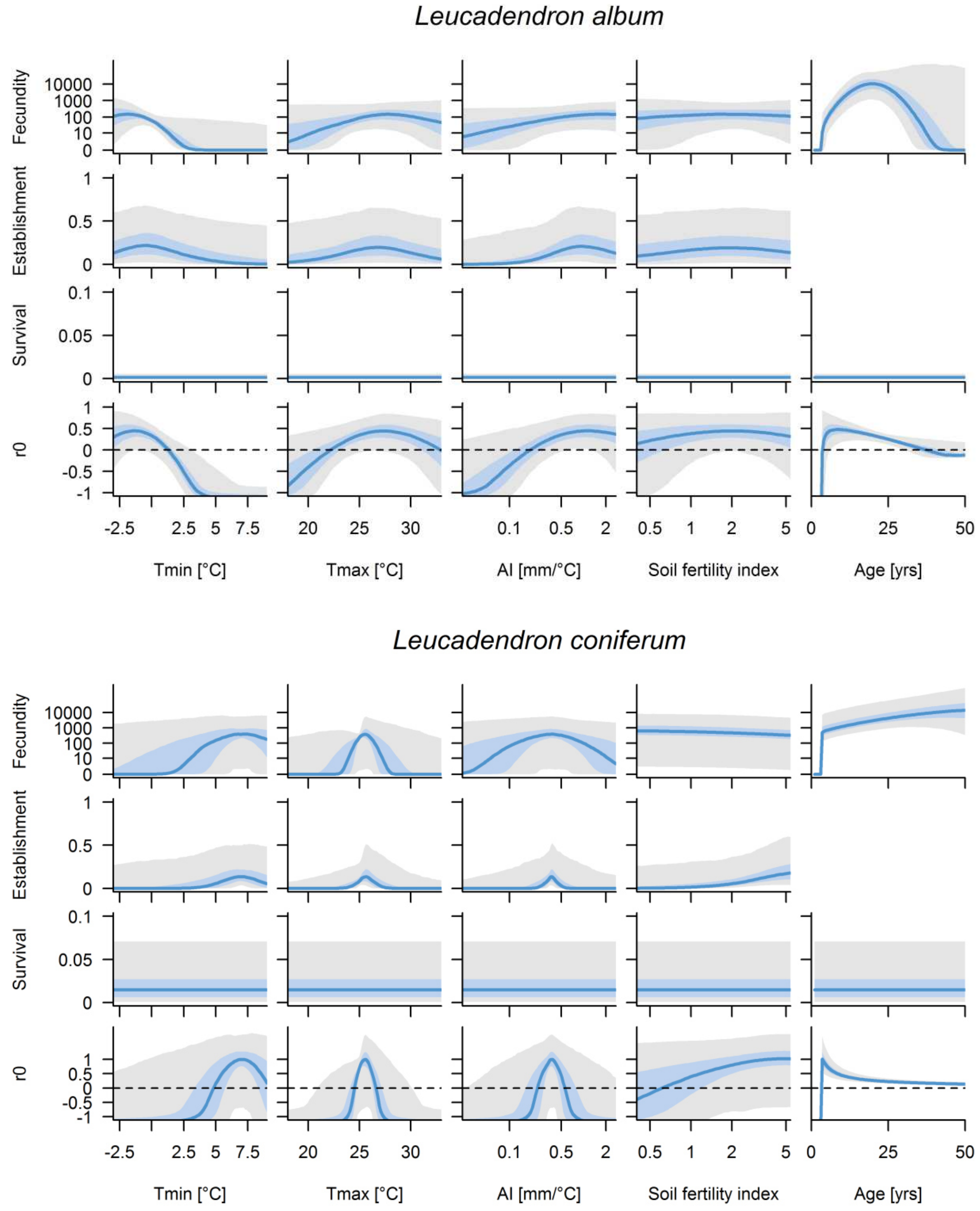


**Fig. S1.** Responses of demographic rates (*μ.fec, π.recr, π.surv*) and intrinsic population growth rate $r_0$ to environmental covariates. Lines show the posterior median of predicted rates and the shaded areas the 50% (dark shading) resp. 95% (light shading) credibility intervals. Response curves were generated by varying each covariate over the range of environmental conditions in the study region while keeping other covariates at the value that optimizes $r_0$.

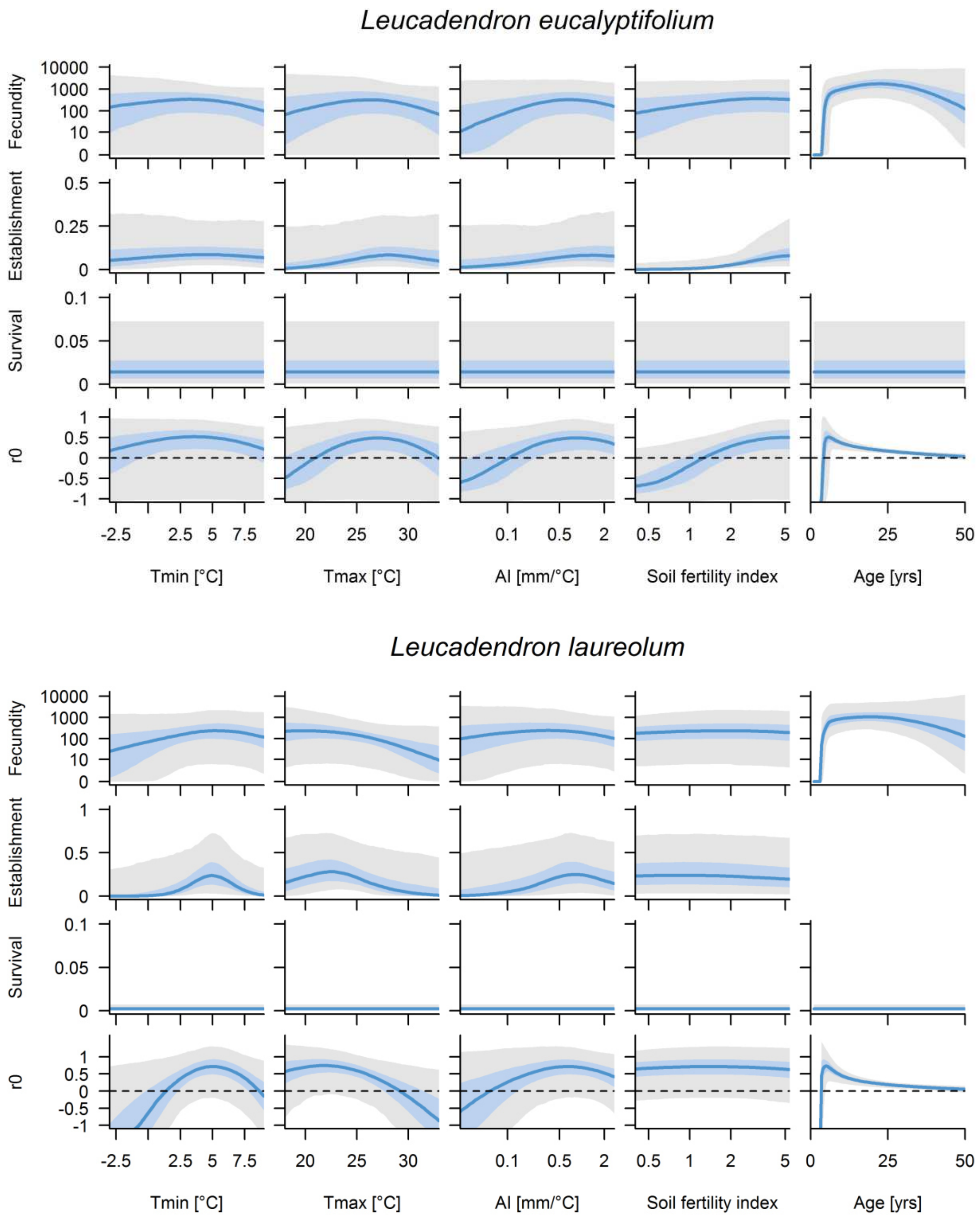


**Fig. S1.** (continued)

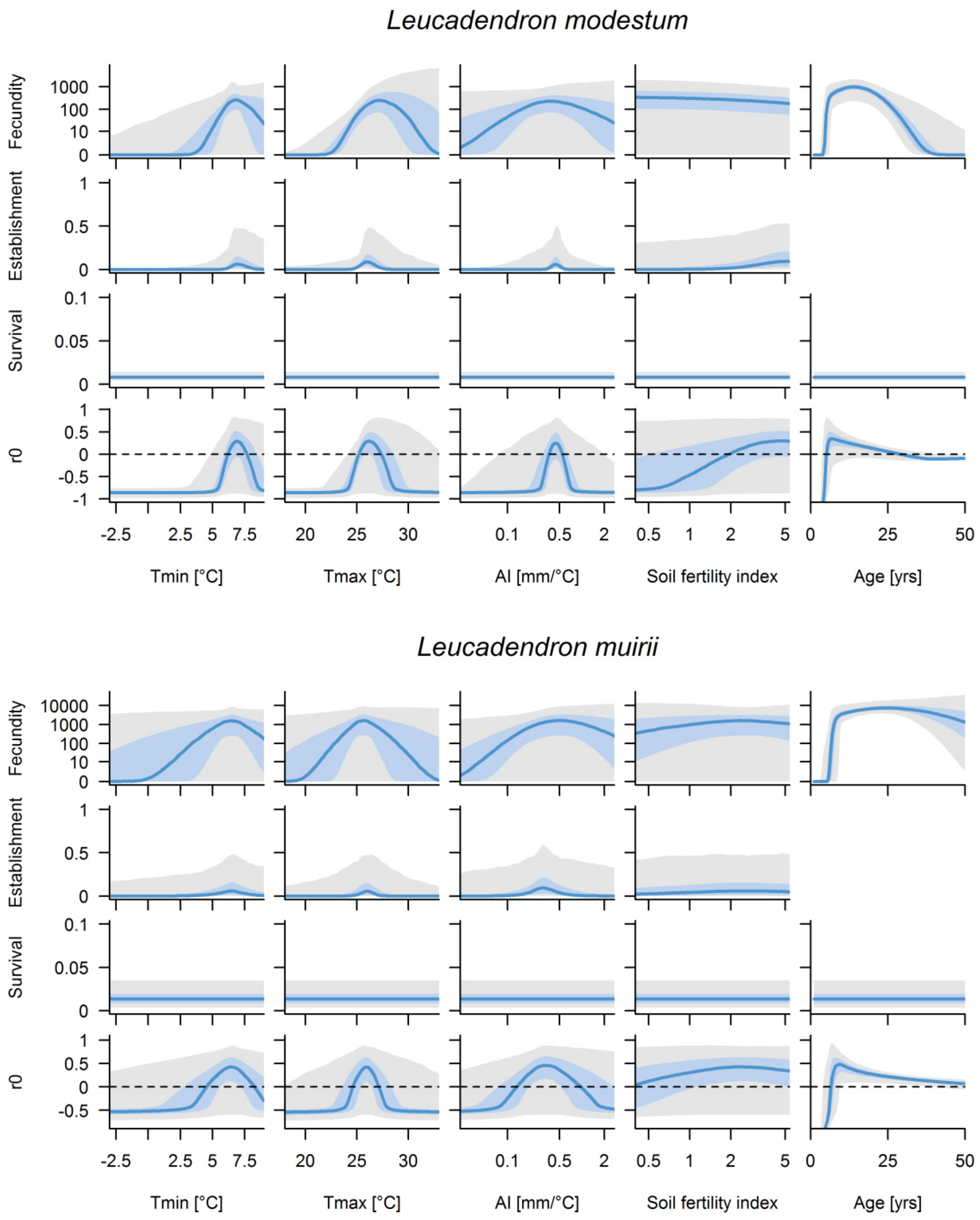


**Fig. S1.** (continued)

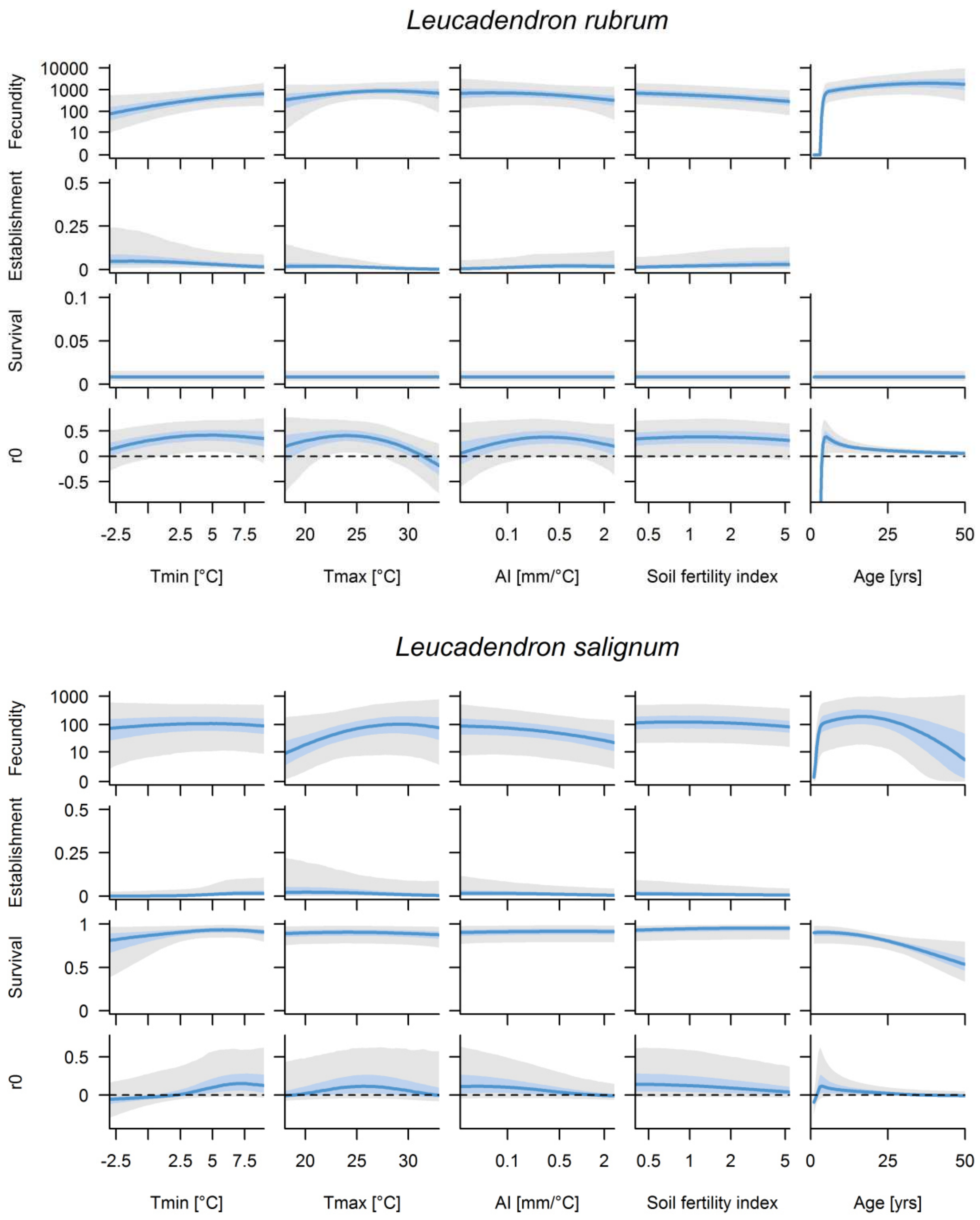


**Fig. S1.** (continued)

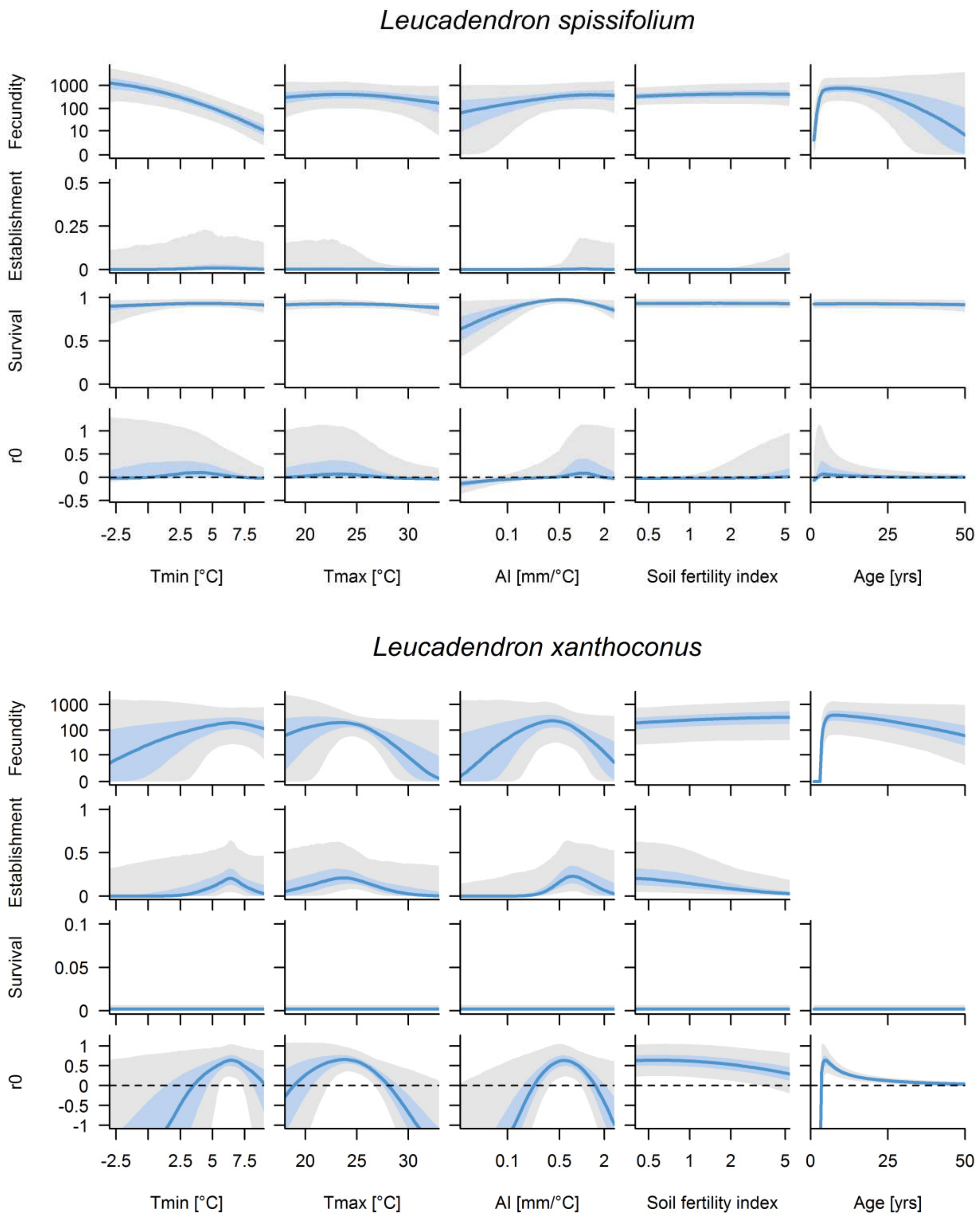


**Fig. S1.** (continued)

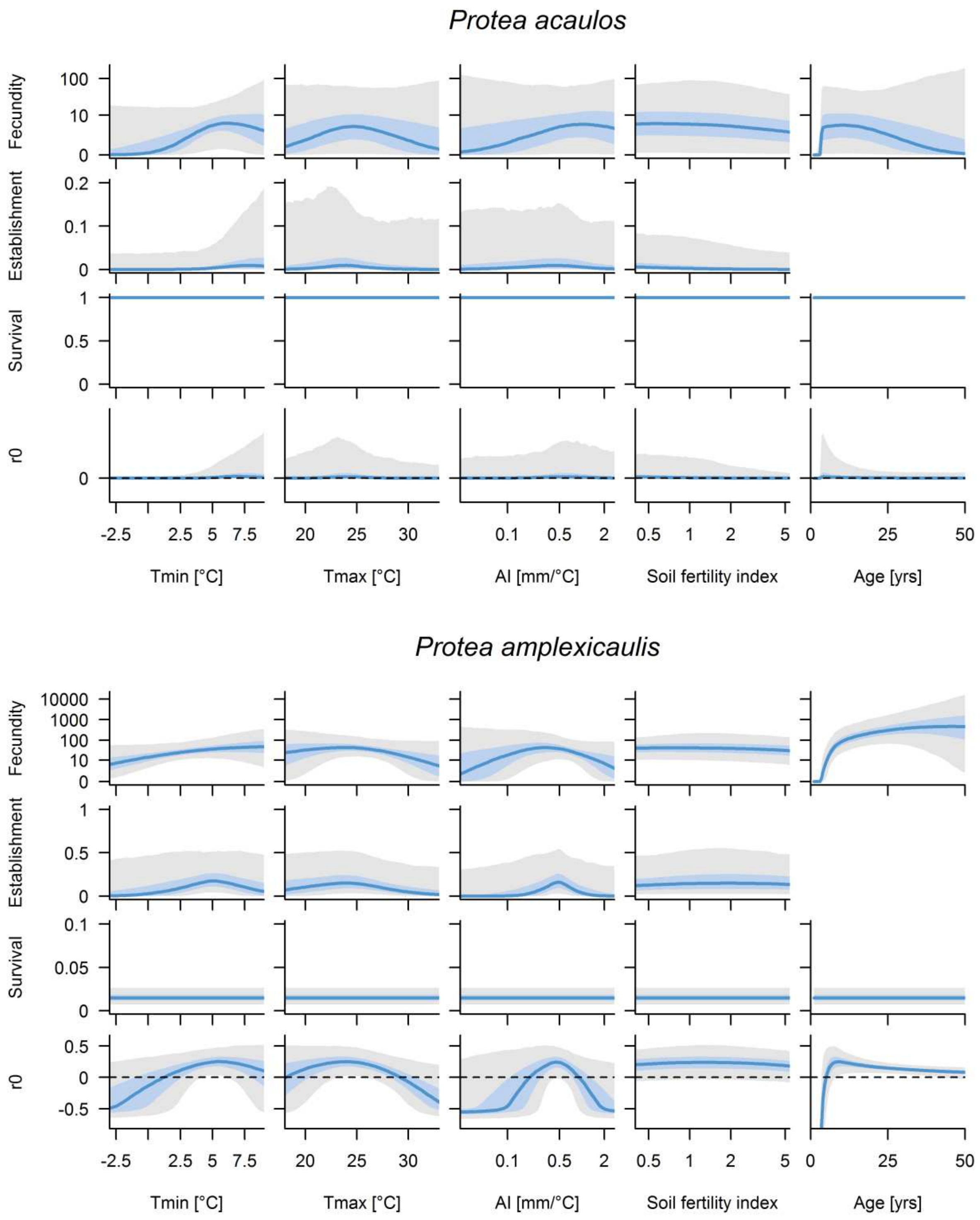


**Fig. S1.** (continued)

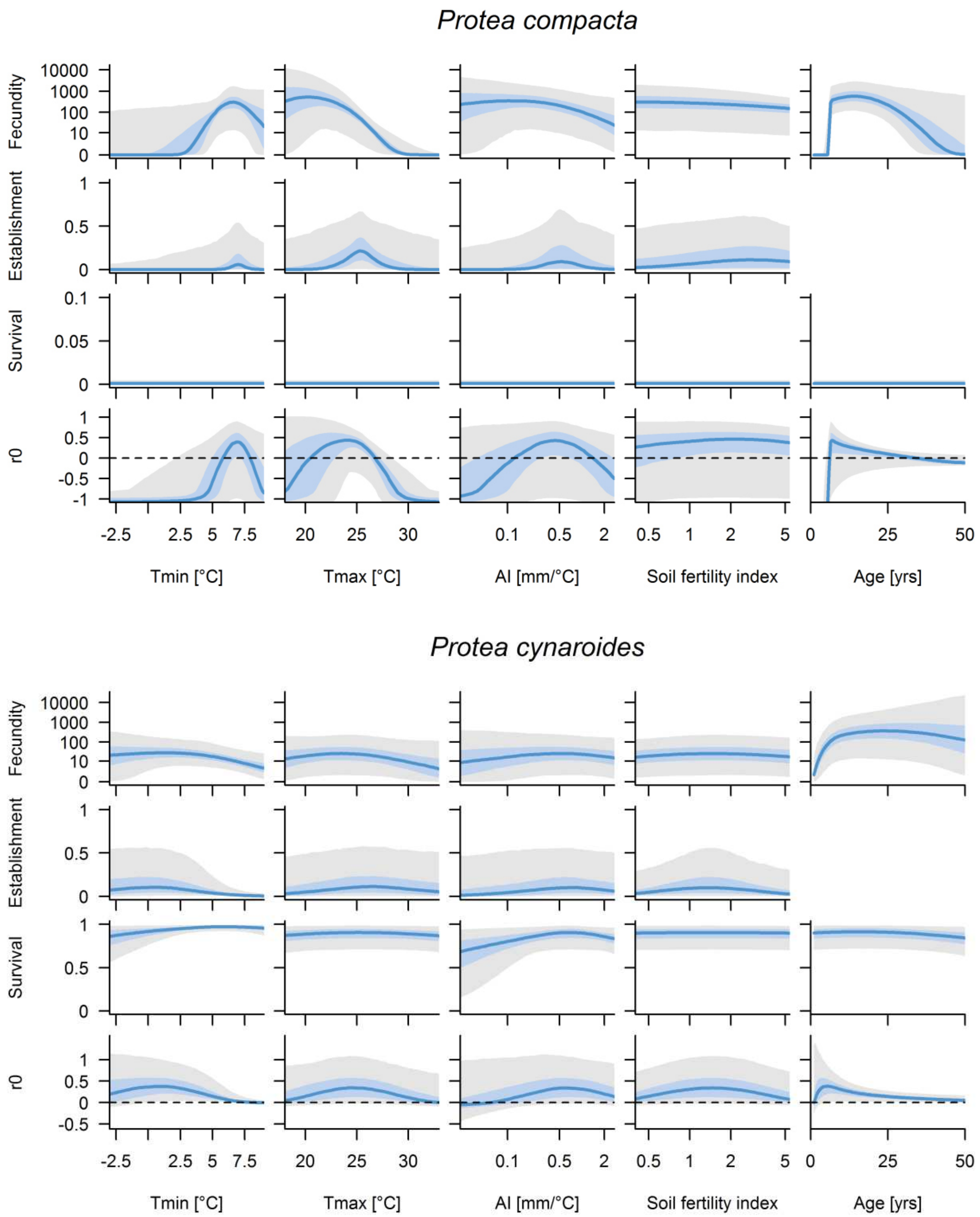


**Fig. S1.** (continued)

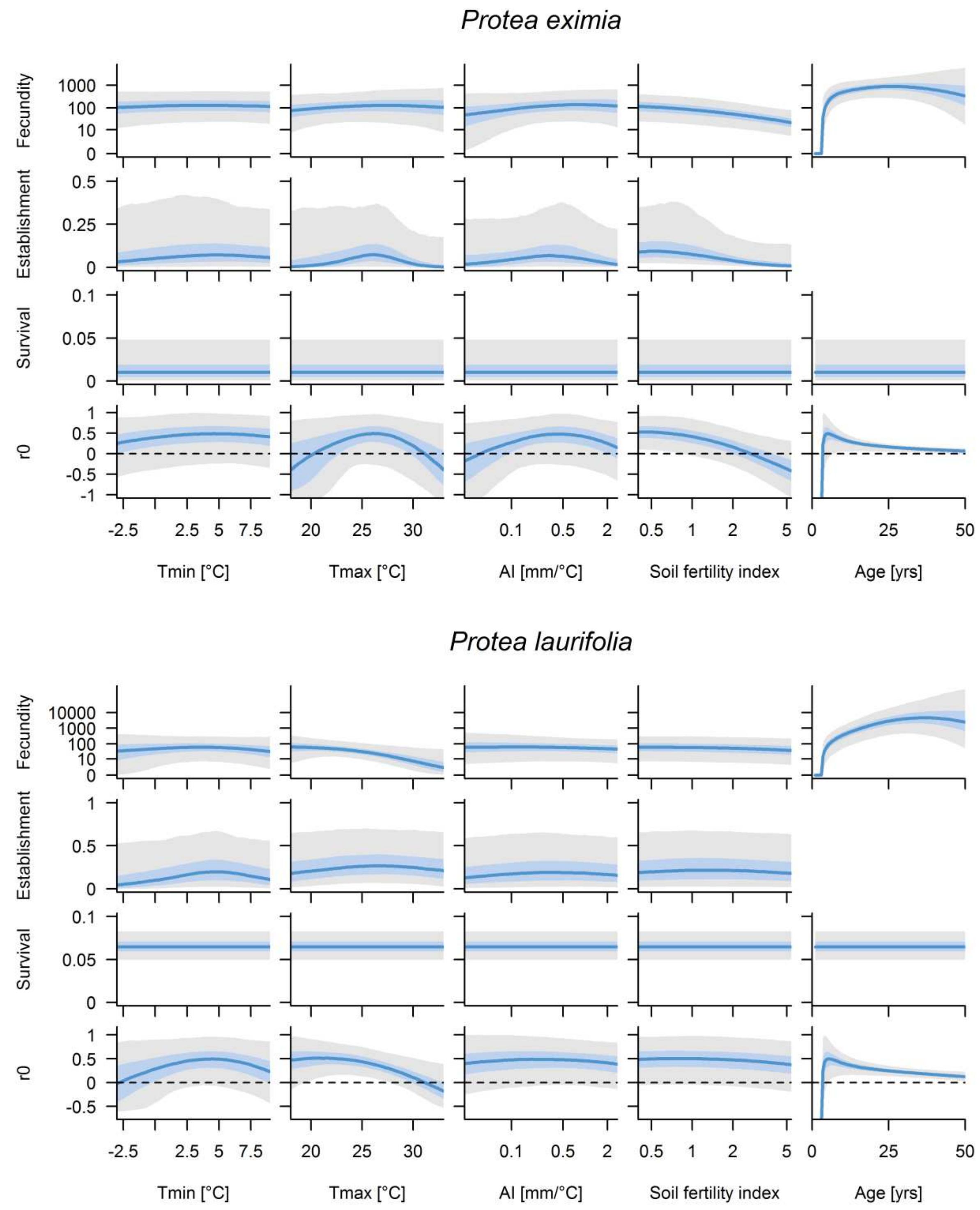


**Fig. S1.** (continued)

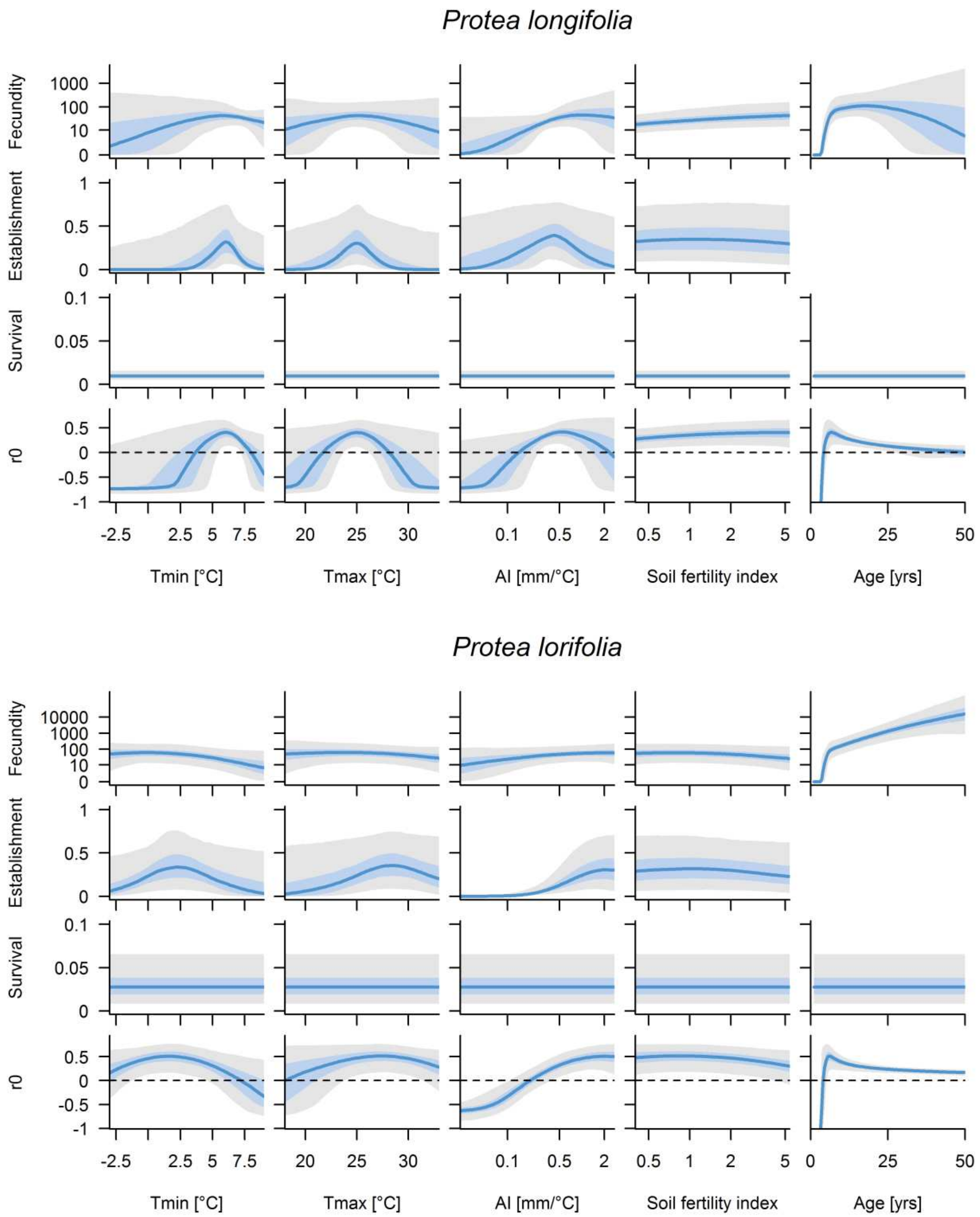


**Fig. S1.** (continued)

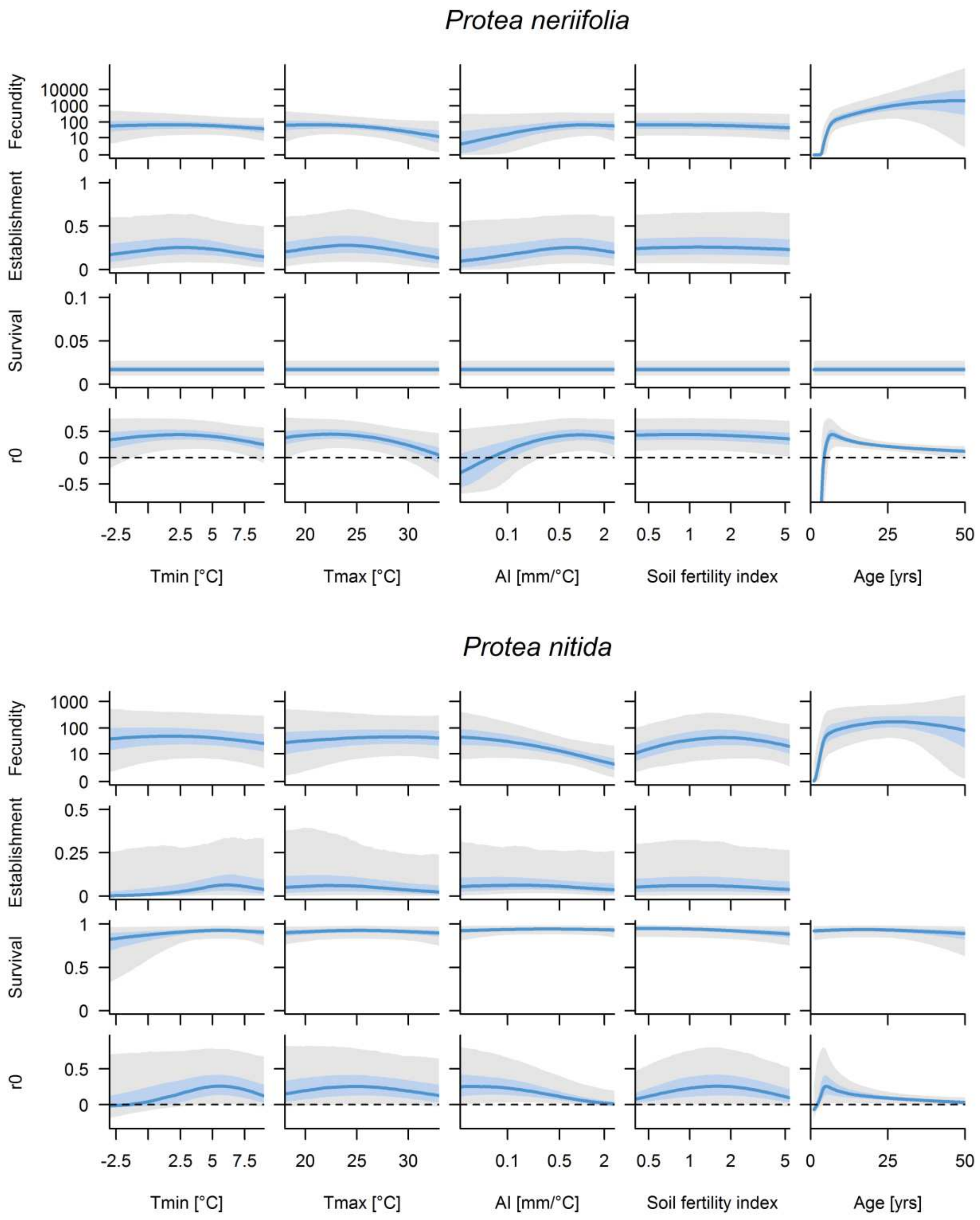


**Fig. S1.** (continued)

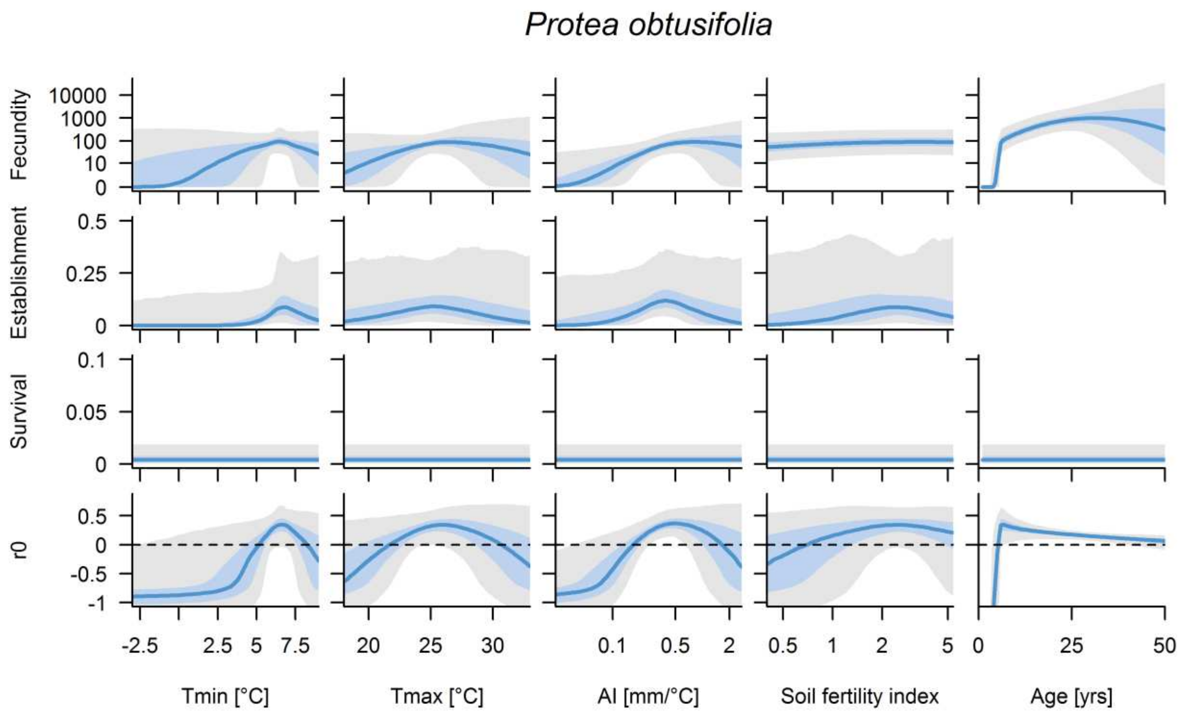


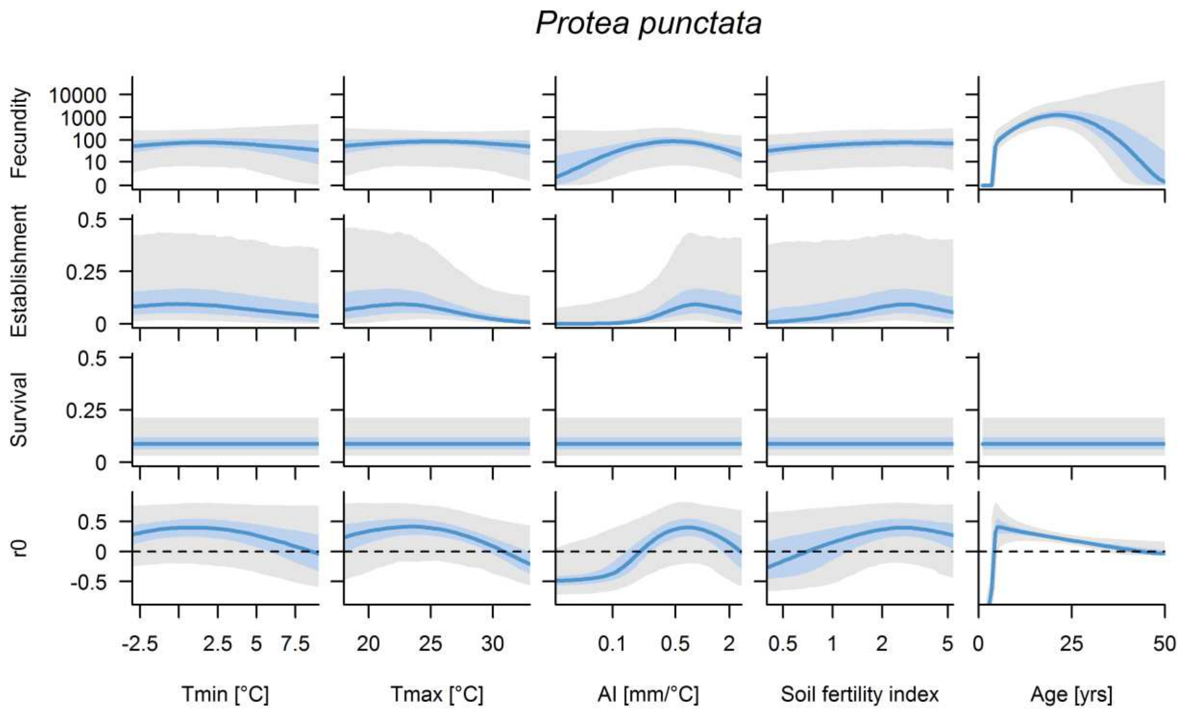


**Fig. S1.** (continued)

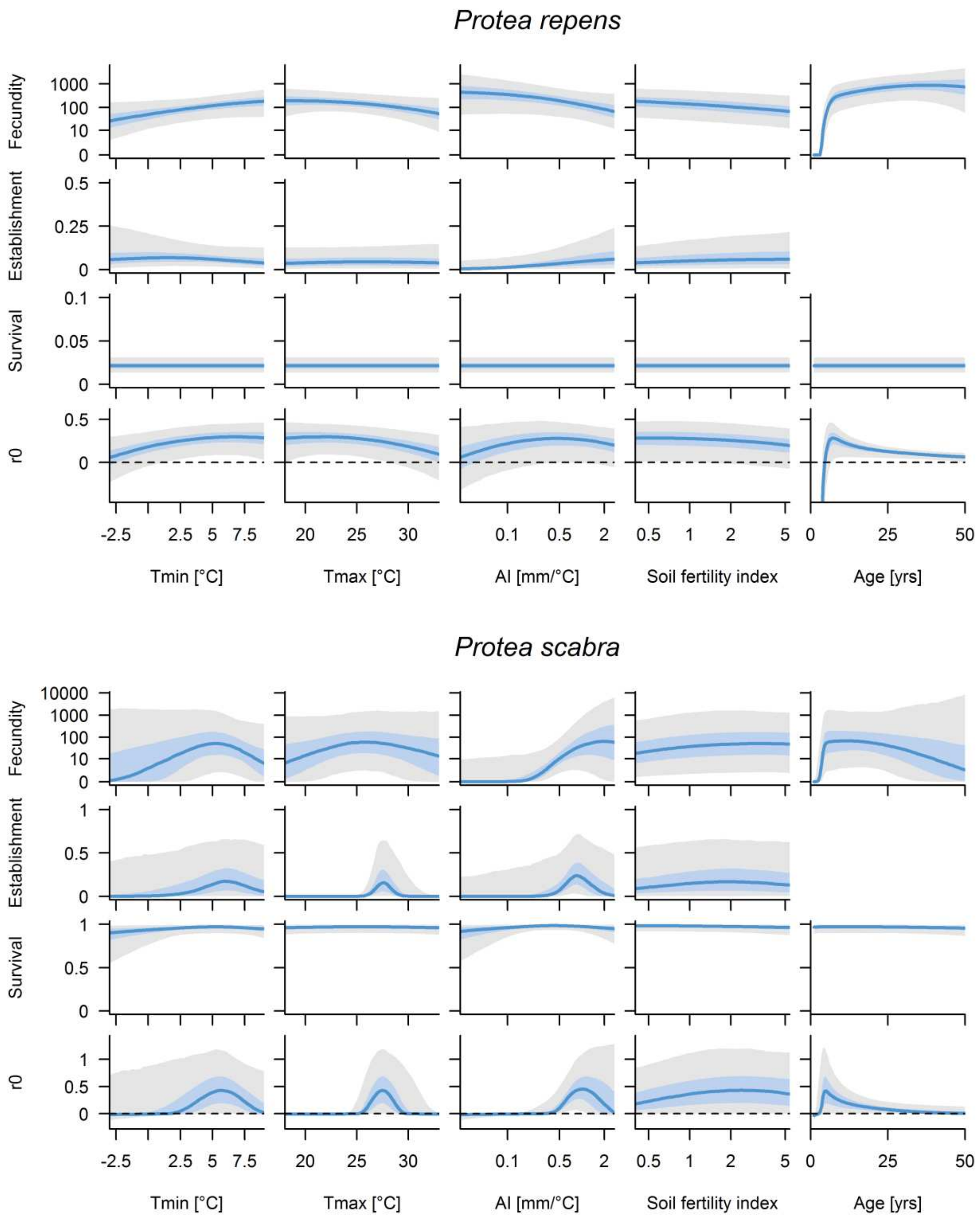


**Fig. S1.** (continued)

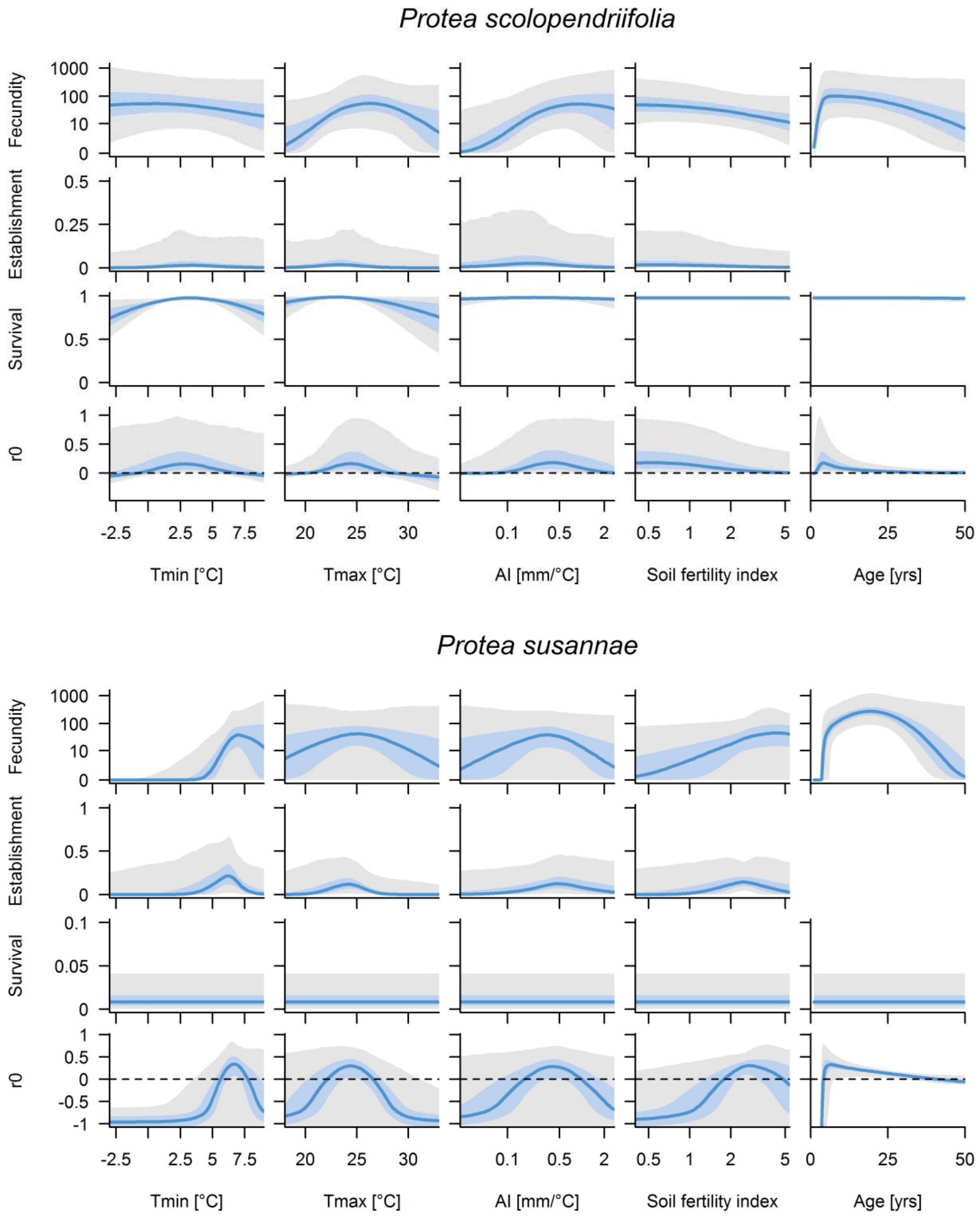


**Fig. S1.** (continued)

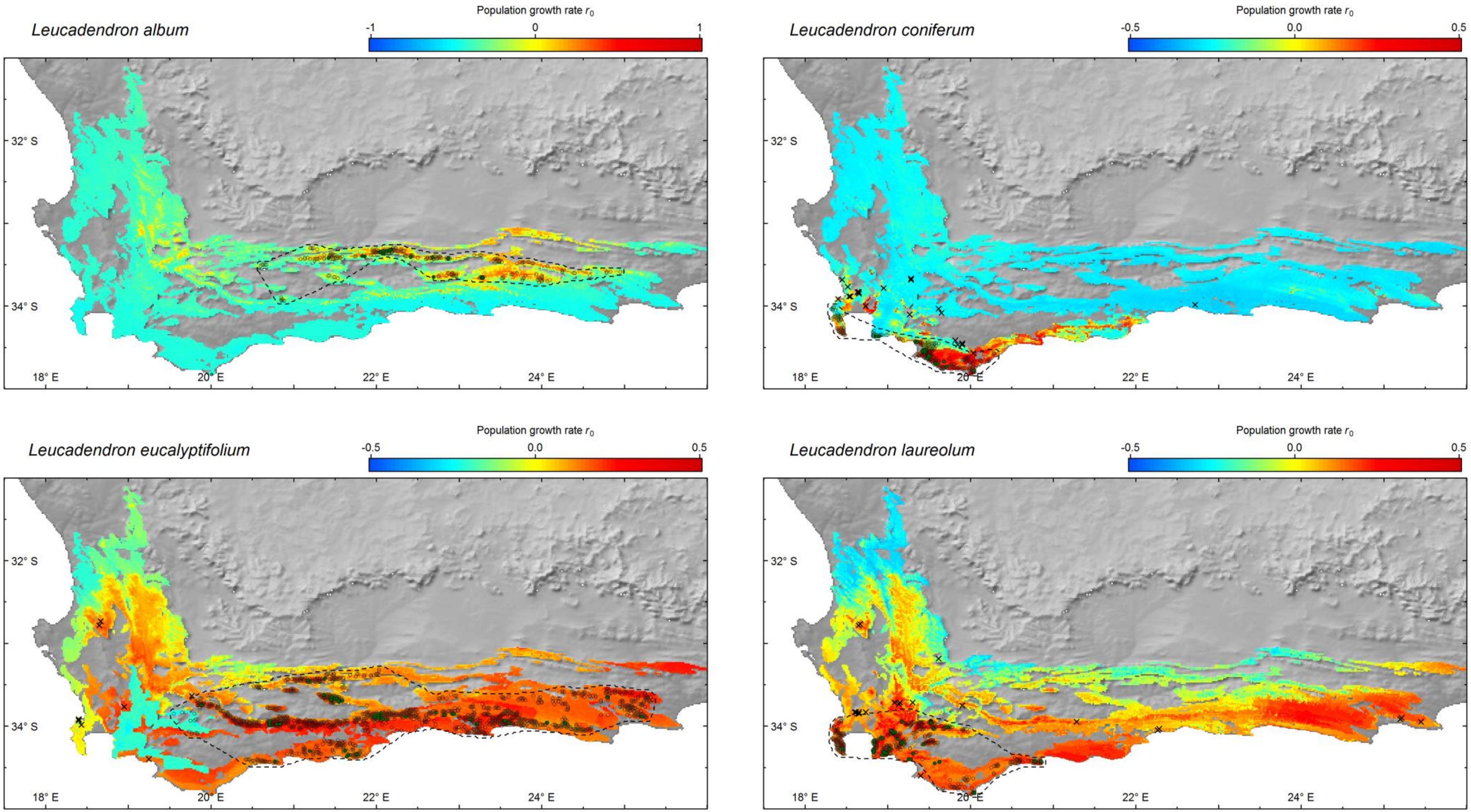


**Fig. S2.** Geographic projection of predicted intrinsic population growth rate $r_0$ across the Fynbos biome (coloured areas) in comparison to the natural geographic range (dashed lines) for each study species. Point symbols show the demographic sampling sites (green circles), presence records of natural populations (open circles) and populations established outside the natural range (crosses). Depicted values of $r_0$ are the medians of the respective Bayesian posterior distributions.

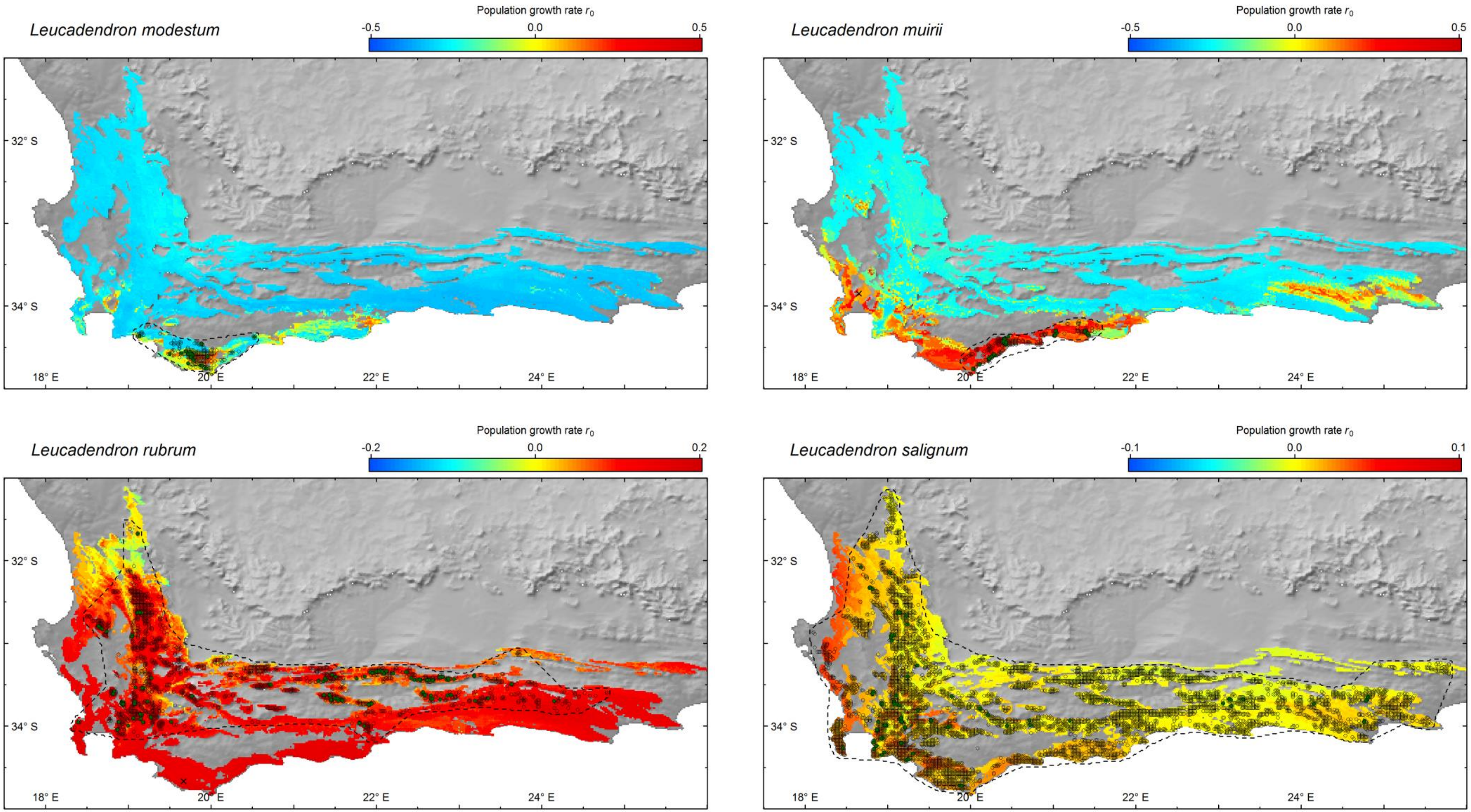


**Fig. S2.** (continued)

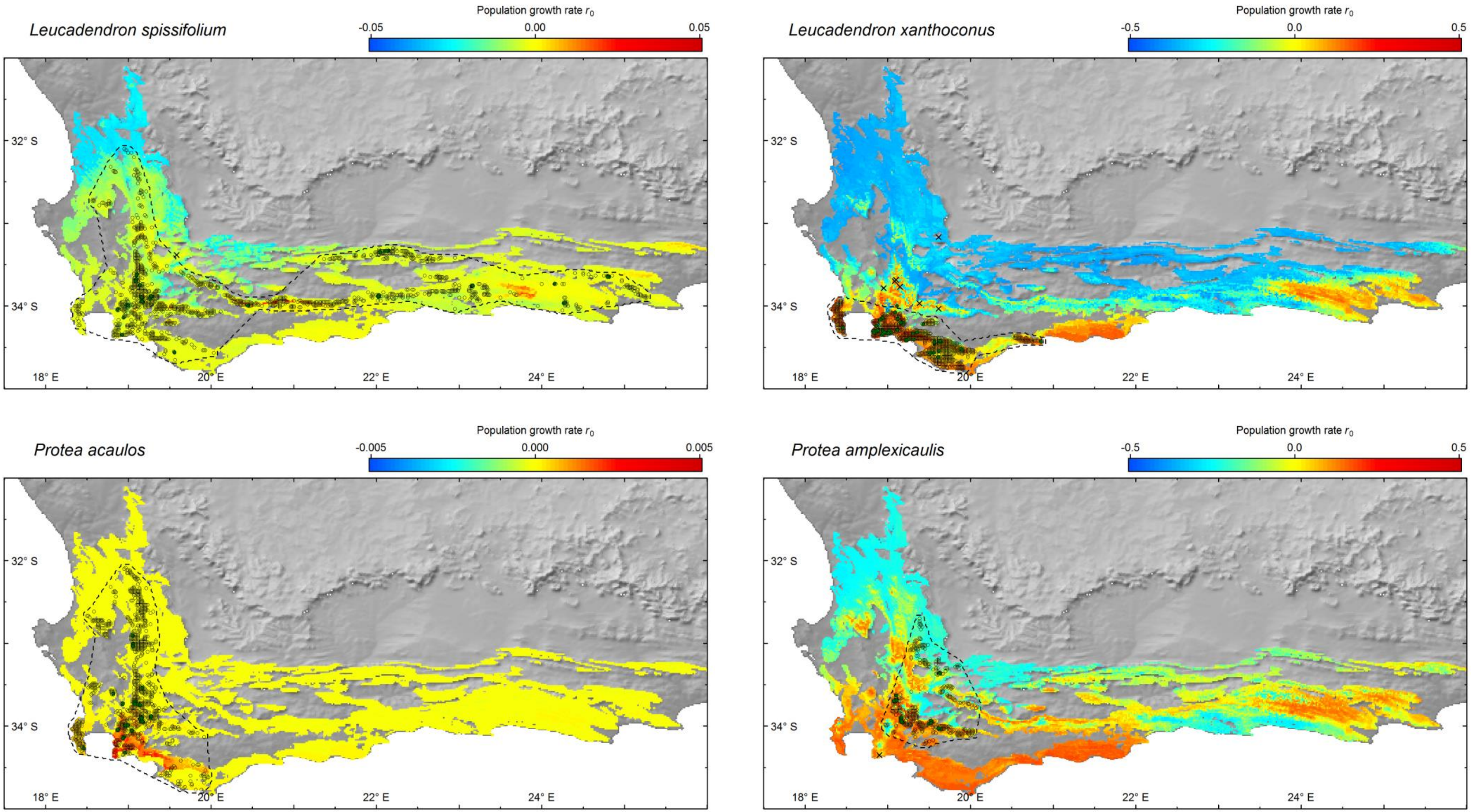


**Fig. S2.** (continued)

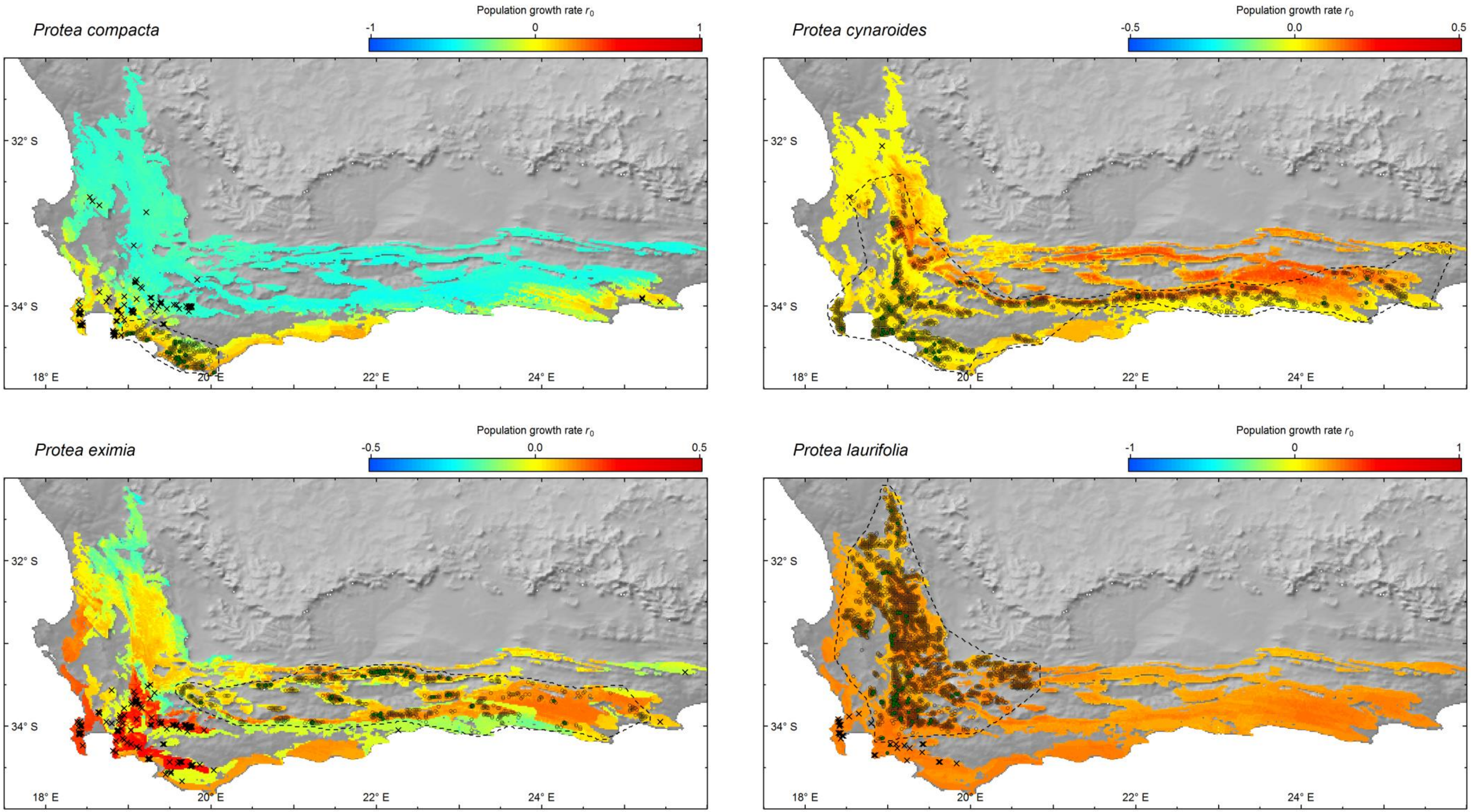


**Fig. S2.** (continued)

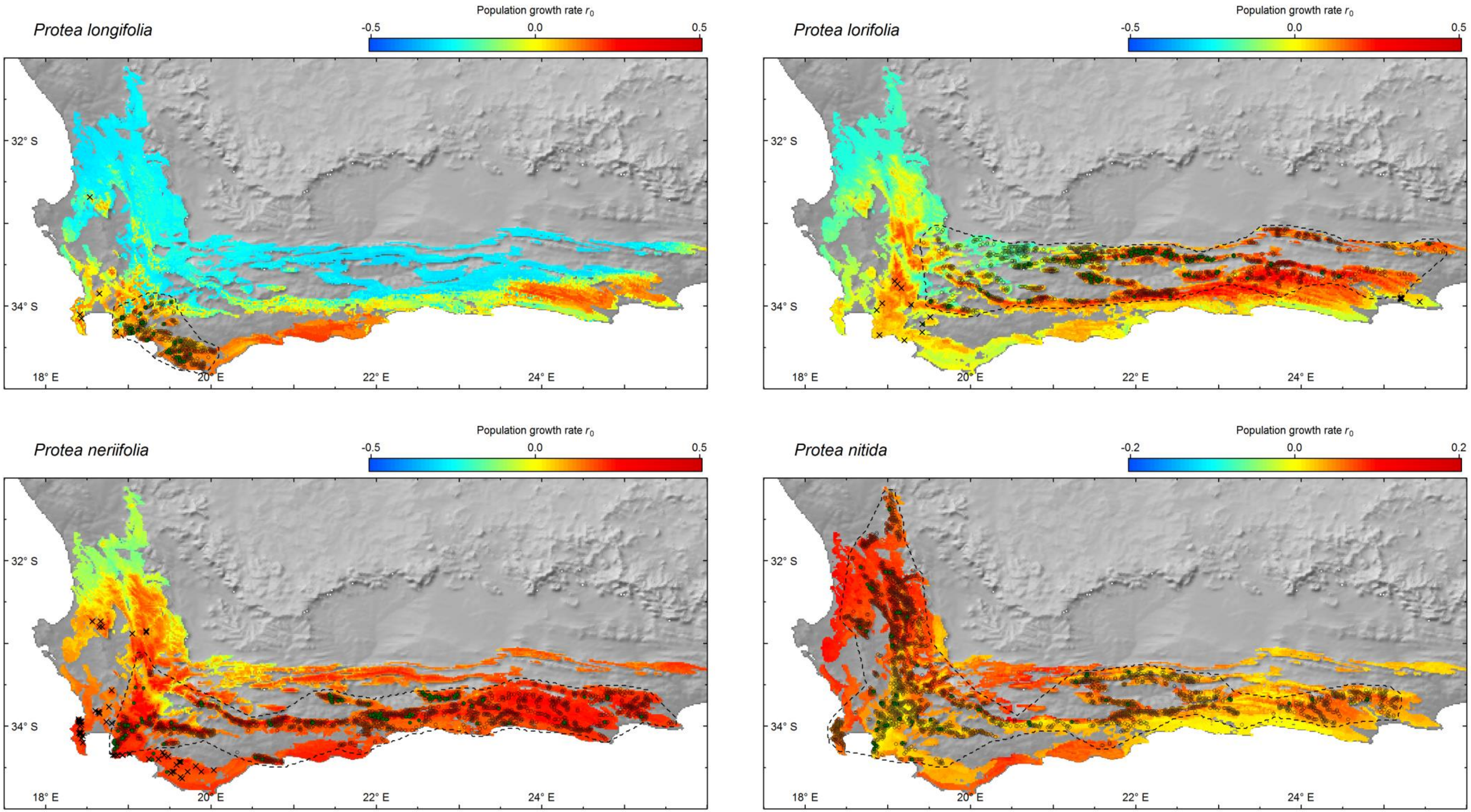


**Fig. S2.** (continued)

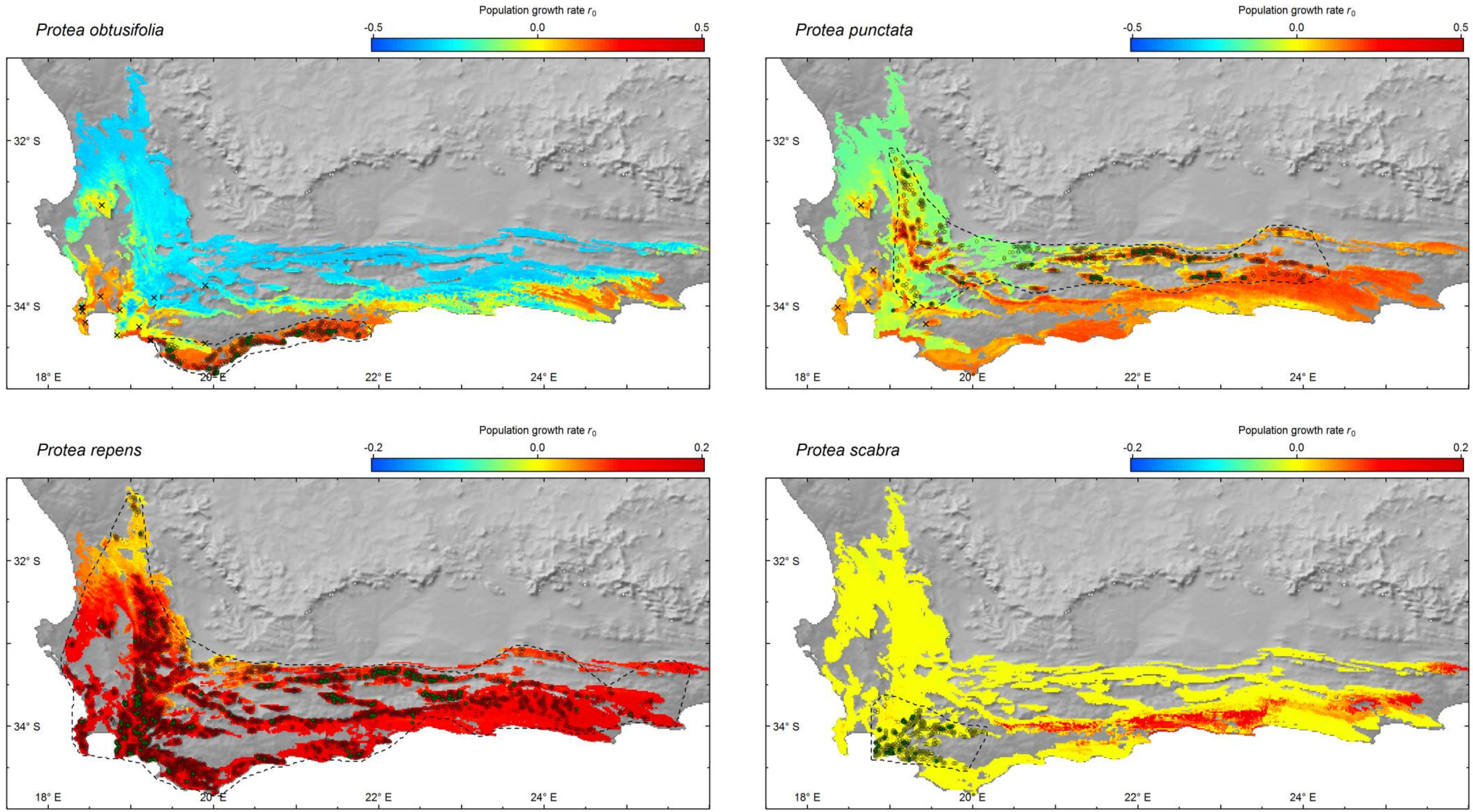


**Fig. S2.** (continued)

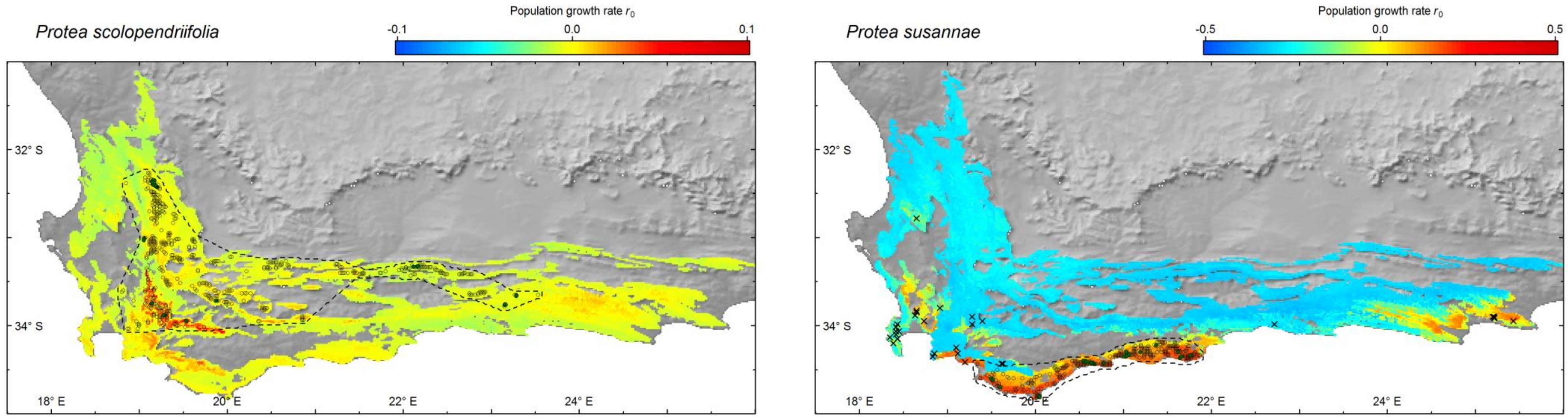


**Fig. S2.** (continued)

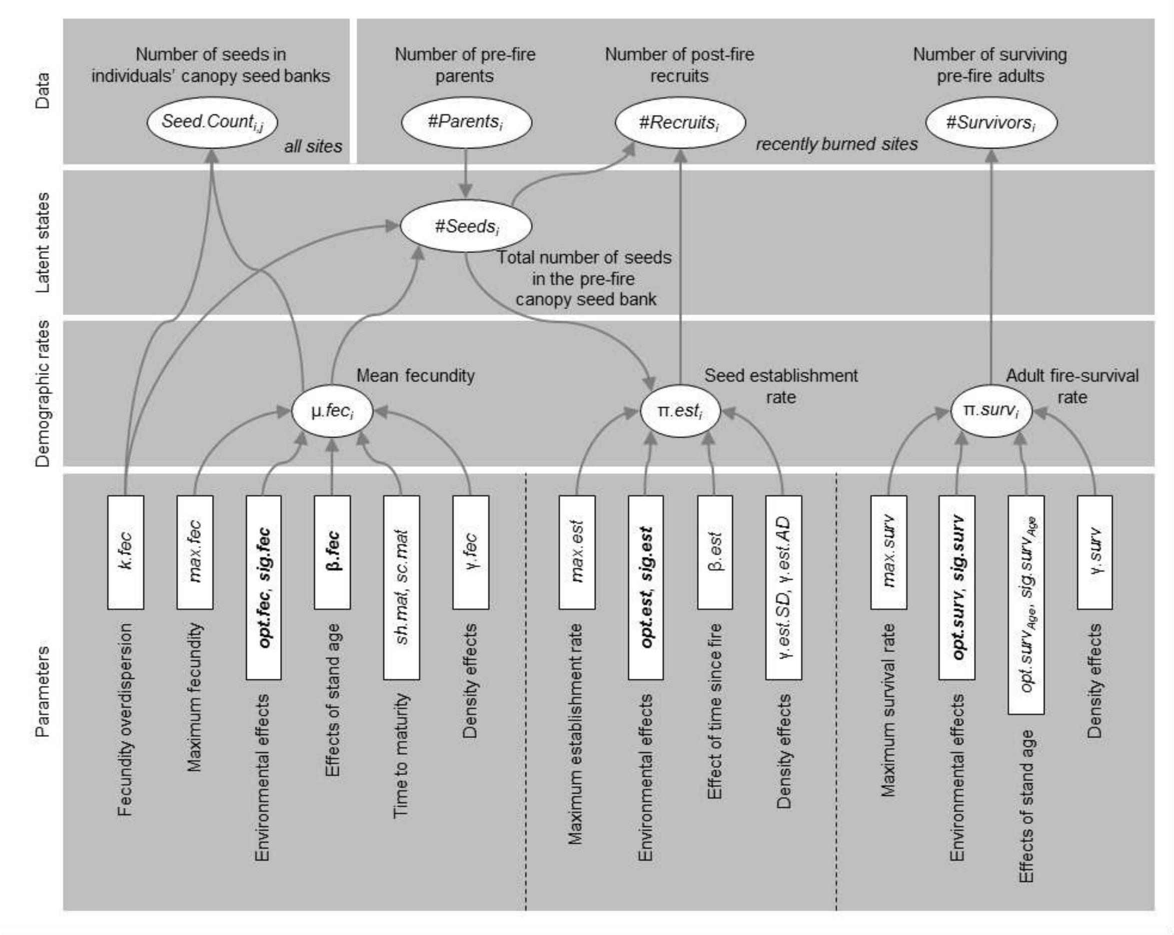


**Fig. S3.** Hierarchical structure of the demographic response model. The model structure is depicted as a directed acyclic graph (DAG) showing the hierarchical relations between model parameters, demographic rates, latent states and the recorded demographic data. The latent state variable #Seeds is included for the inference of per-seed establishment rates from the observed number of recruits on recently burned sites. On these sites the number of pre-fire parents can be determined by combining counts of burned skeletons and fire-surviving adults, but no data on the pre-fire canopy seed bank is available. Hence the size of the pre-fire canopy seed bank on these sites is predicted from the fecundity submodel. See the Methods for a detailed description of all variables and their statistical relations and Tab. S5 for prior distributions of the model parameters.

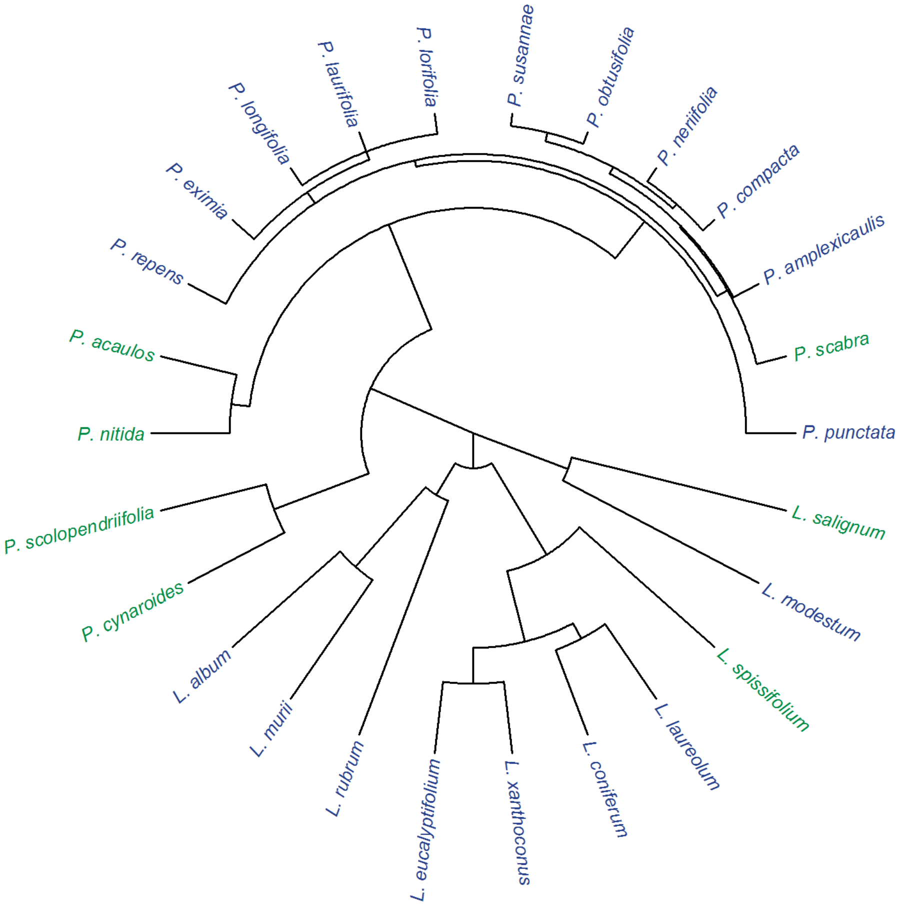


**Fig. S4.** Phylogenetic tree of the study species. The phylogenetic reconstruction is based on 18 DNA markers for 291 taxa of the Proteaceae family. Here only the trimmed phylogenetic tree for the 26 study species is shown (blue: nonsprouters, green: resprouters).
